# Genomic divergence of sympatric lineages within *Stichopus* cf. *horrens* (Echinodermata: Stichopodidae): Insights on reproductive isolation inferred from SNP markers

**DOI:** 10.1101/2024.10.29.620868

**Authors:** Kenneth M. Kim, Apollo Marco D. Lizano, Robert J. Toonen, Rachel Ravago-Gotanco

**Affiliations:** Department of Ecology and Evolution, University of Lausanne, Lausanne, Switzerland; Swiss Institute of Bioinformatics, Lausanne, Switzerland; Marine Science Institute, University of the Philippines, 1101, Diliman Quezon City, Philippines; Hawai‘i Institute of Marine Biology, School of Ocean and Earth Science and Technology, University of Hawai‘i at Mānoa, Kāne‘ohe, HI 96744, USA

## Abstract

How reproductive barriers arise in early stages of divergence among broadcast spawning organisms that exist in sympatry remains poorly understood. Reproductively isolated lineages (Clade A and B) of *Stichopus* cf. *horrens* were previously reported across the western Pacific, with an additional putative cryptic species detected within Clade B lineage warranting further examination. The present study further examines the hypothesis that the two mitochondrial lineages (Clade A and Clade B) of *Stichopus* cf*. horrens* represent putative cryptic species and whether another cryptic species within the Clade B lineage exists using a reduced representation genomic approach. Using double-digest RAD (ddRAD) sequencing, a total of 9,788 single nucleotide polymorphism (SNP) markers were used to examine divergence among *Stichopus* cf*. horrens* lineages (n = 82). Individuals grouped into three SNP genotype clusters, broadly concordant with their mitochondrial lineages and microsatellite genotype clusters, with limited gene flow inferred among clusters. Outlier analysis recovered highly divergent SNP loci with significant homology to proteins related to rhodopsin and tachykinin receptor signaling, sperm motility, transmembrane transport and hormone response. This study confirms the existence of three reproductively isolated genotype clusters within *Stichopus* cf. *horrens* and highlights gene regions related to reproduction that may contribute to establishing reproductive barriers between broadcast spawning species at an early stage of divergence.

## 1 Introduction

With the use of molecular approaches for species delimitation, cryptic species are now more readily detected and appear to be common in the marine environment (Bickford et al. 2007; Knowlton 1993, 2000; Nygren, 2014). While allopatric speciation can be easily explained by physical barriers when cryptic species have disconnected distribution ranges, species with parapatric, overlapping, or sympatric ranges likely have a more complex evolutionary history (Bird et al., 2012; Faria et al., 2021). This complexity is often seen in the marine environment, where barriers to gene flow are less obvious and divergences along ecological boundaries are thought to occur more frequently than in the terrestrial environment (Bowen et al. 2013). In the marine environment, where external fertilization is common, it is more challenging for extrinsic ecological barriers to arise, leading to increased opportunities for hybridization and decreased opportunities for divergence. Mechanisms like gametic incompatibilities and variations in the timing and spatial variation of gamete release are considered key factors maintaining reproductive isolation in broadcast spawning species occurring in sympatry (Palumbi 1994, Swanson & Vacquier, 2011; Bird et al., 2011). The evolution of these mechanisms to limit hybridization are acquired in various orders throughout the “speciation continuum” (De Queiroz, 2005, 2007; Kulmuni *et al*., 2020; Nosil and Feder 2012; Seehausen *et al.,* 2014; Roux *et al*., 2016; Stankowski and Ravinet 2021) and the existence of cryptic species offer an interesting toolbox to examine how reproductive barriers arise in early stages of divergence among broadcast spawning organisms that exist in sympatry.

The tropical sea cucumber *Stichopus* cf*. horrens*, exhibits a high degree of morphological variability making species identification challenging. A recent phylogenetic study revealed two divergent lineages (Clade A and B) within *S.* cf. *horrens* where identification based on molecular data are incongruent with spicule morphology (Lizano et al., 2024). These lineages were further revealed to be reproductively isolated, with limited contemporary gene flow among sympatric populations. Moreover, an additional putative cryptic species within the Clade B lineage was also detected warranting further examination. The low frequency of mitonuclear discordance coupled with the absence of F1 hybrids between lineages raises the question: how is reproductive isolation maintained between broadcast spawning cryptic species despite occurring in sympatry? Prezygotic isolation, particularly through the accumulation of gamete incompatibilities due to selection, is thought to be a primary mechanism reinforcing isolation in broadcast spawning marine organisms in sympatry (Levitan et al. 2004; Metz and Palumbi 1996; Landry et al., 2003; Coyne and Orr, 2004; Bird et al., 2012). Positive selection on gamete recognition proteins (GRPs) has been reported in several echinoderm species (Lessios 2011; Sunday and Hart 2013; Patiño et al., 2016) and these proteins have been shown to establish reproductive barriers between closely related broadcast spawning individuals in sympatry (Palumbi 2009; Vacquier and Swanson 2011; Kosman and Levitan 2014). The role of GRPs in forming prezygotic reproductive barriers in sea cucumbers, however, is not yet known, and it remains unclear whether divergence in GRPs contributes similarly to speciation in other broadcast spawning organisms. Levels of polymorphism and patterns of divergence in GRPs vary across different groups. For example, in *Ciona intestinalis*, although GRPs are found to evolve more rapidly than proteins with other functions, the rates of evolution between allopatric and sympatric populations of the two reproductively isolated forms were not significantly different (Nydam and Harrison 2011). Similarly, in the sea stars *Cryptasterina hystera* and *C. pentagona,* little evidence of selection on gamete recognition genes was observed. Instead, signatures of selection were found in genes related to environmental interactions (Guerra et al., 2021). In addition, in *Ophioderma longicauda*, there is no evidence of positive selection on gamete recognition proteins, rather, positive selection was found on two genes encoding sperm ion channels involved in signal transduction cascade in response to pheromones (Weber et al. 2017). Alternative mechanisms, such as habitat preference and synchronized spawning triggered by hormonal cues (references: Bierne et al., 2003; Hamel and Mercier, 1995, 1999; Mercier and Hamel, 2010), have been proposed. This implies that other genes involved in reproduction, potentially those upstream in the reproductive cascade, may also play a role in prezygotic isolation in sympatric broadcast spawning species.

In this study, we utilized single nucleotide polymorphisms (SNPs) analyzed via double-digest restriction-site-associated DNA sequencing (ddRADseq; Peterson et al., 2012) to test whether similar patterns of genetic differentiation and clustering previously found within *Stichopus* cf. *horrens* using mitochondrial and microsatellite markers will be recovered. Specifically, we expect genotype clusters of *Stichopus* cf. *horrens* recovered using SNP loci to be concordant with mitochondrial lineages and microsatellite genotype clusters, and further exhibit significant genetic differentiation resulting from limited gene flow. Additionally, this study will use SNP outlier analysis to examine the hypothesis that higher levels of genetic divergence will be detected in gene regions or loci related to reproduction (e.g. GRPs). Genome scans are now commonly used to identify potential outlier loci by examining molecular signatures of selection (Ravinet et al., 2017). These techniques involve assessing genetic divergence (e.g., based on distance metrics such as *FST* or *dxy*) or admixture proportions (Martin et al., 2015) across thousands to millions of loci to pinpoint regions of the genome where variation deviates from neutral divergence models.

The previous study reported the existence of two cryptic species and suggested an additional third species but with weak support due to small sample sizes and limited resolution from 6 microsatellites markers (Lizano et al., 2024). Here in this study, using high resolution SNP markers, we recovered three genotype clusters exhibiting limited gene flow between them. This results provide evidence of the existence of three cryptic species within *Stichopus cf. horrens*.

In addition, we did not recover highly divergent SNP loci with significant homology to GRPs instead we found other proteins that are related to receptor signaling, neurotransmitter receptor activity, response to testosterone, peptide hormone processing and sperm motility. Findings from this study will provide valuable insights into the possible drivers of reproductive isolation in broadcast spawning organisms occurring in sympatry.

## 2 Materials and Methods

### 2.1 Sample collection and DNA extraction

This study used *Stichopus* cf. *horrens* individuals collected from three sites in the Philippines where cryptic species have been reported and occur in sympatry (Lizano et al. 2024): Sta. Ana, Cagayan (STA) and two sites in the Davao Gulf: Mabini, Davao de Oro (MAB), and Pujada Bay, Davao Oriental (PUJ) (Figure 1). Genomic DNA was extracted from preserved tissues (body wall or papillae in 96% Ethanol or RNAShield^TM^) using a QIAGEN DNEasy Blood and Tissue kit. DNA concentration was quantified using a fluorometer, and quality assessed by agarose gel electrophoresis. Of the 96 samples used in this study, some samples have been previously genotyped at the mitochondrial COI region (n = 69) and at 6 microsatellite loci (n = 55) for assignment to mitochondrial lineage and genotype clusters (Lizano et al. 2024, Table S1).

**Figure 1.**
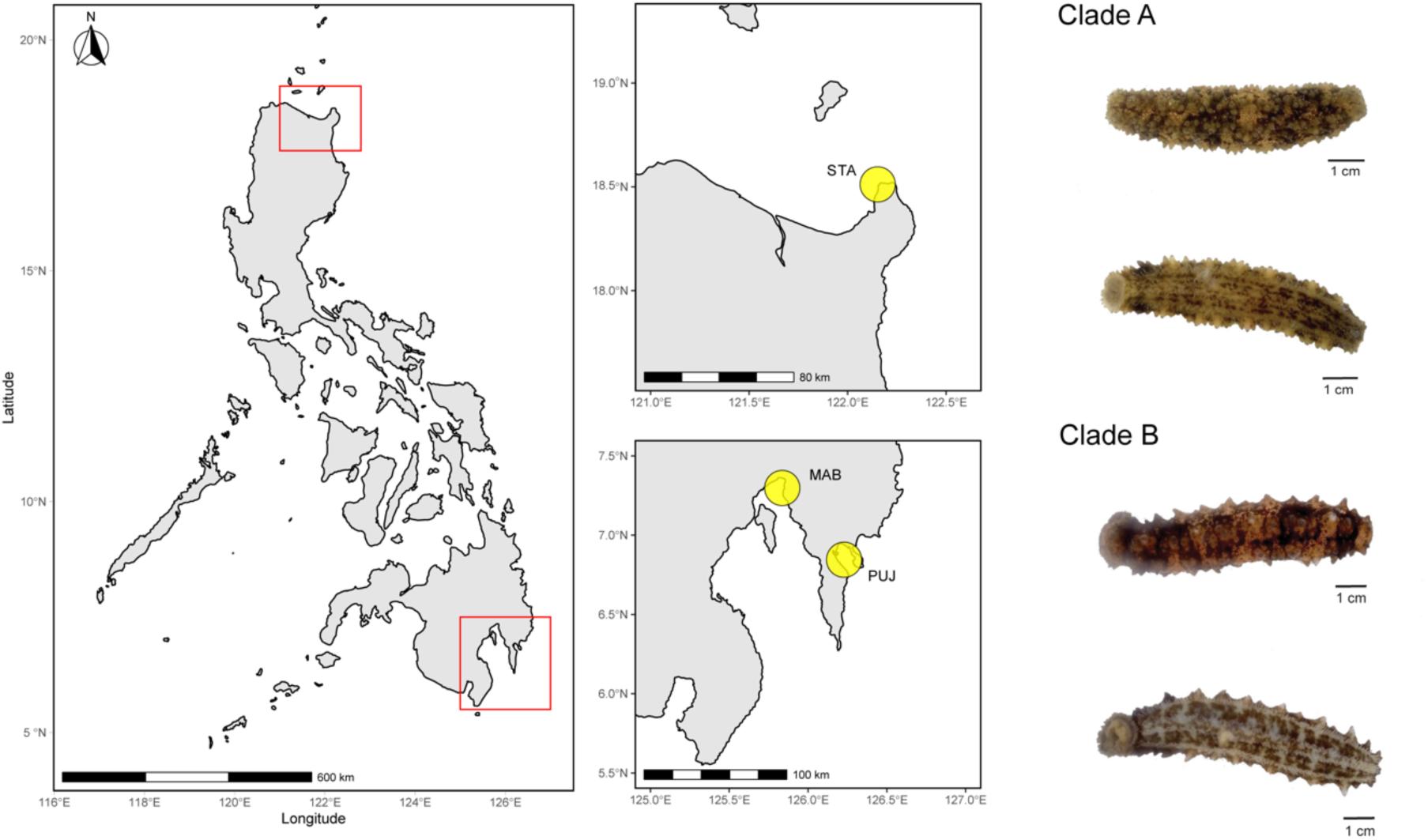
Sampling locations for *Stichopus* cf. *horrens*. Samples were collected from three sites in the Philippines (left) where three putative pseudocryptic species were reported to be sympatric: Sta. Ana, Cagayan (STA) (top middle), Mabini, Davao de Oro (MAB), and Pujada Bay, Davao Oriental (PUJ) (bottom middle). Photographs of two cryptic species within *S.* cf. *horrens*, designated as Clade A and Clade B (Lizano et al. 2024) are shown (right). Maps were generated in R using the sf, naturalearth, and ggplot2 packages.

### 2.2 Double-digest restriction associated DNA sequencing and quality filtering

Library preparation and sequencing were performed at the Genomics Core Lab, Texas A&M University Corpus Christi. Libraries were prepared following the double digest RADseq (ddRADseq) protocol adapted from Peterson et al. (2012) using restriction enzymes *PaeI* and *TasI* for digestion of genomic DNA. Individually barcoded libraries were pooled, size-selected, and sequenced on one lane of Illumina HiSeq 4000 (paired-end, 150 bp). Demultiplexed raw reads were quality assessed using FASTQC (Andrews 2010). Reads were initially filtered for quality using the ‘process_radtags.pl’ pipeline in STACKS version 1.48 (Catchen et al. 2011). Individual reads with phred scores below 10 or with ambiguous barcodes were discarded. Quality filtered reads were then filtered for contaminants using BBMap and BBSplit (https://github.com/BioInfoTools/BBMap) by mapping against bacterial and viroid sequences.

### 2.3 De novo assembly and SNP filtering

RADseq loci were de novo assembled using the ‘denovo_map.pl’ pipeline in STACKS. Different parameters were explored and evaluated based on the number of generated loci and the depth of coverage per sample. The final parameters selected for this study that yielded a suitable number of loci and minimum depth of coverage (10x) were a minimum read depth to create a stack (-m = 3), number of mismatches allowed between loci within individuals (-M = 7), and number of mismatches allowed between loci within catalog (-n = 8). The parameters used followed recommendations by Paris et al. (2017) where -*M* should be high for populations with high divergence and *-n* is either =*M*, *M-1* or *M+1*.

After de novo mapping, the ‘populations’ module of STACKS was used to filter loci. We used a relaxed filtering criterion, retaining SNPs shared by 60% percent of individuals across all populations (-r = 60), retaining only one SNP per locus (-write_single_snp). SNPs with a minor allele frequency (MAF) less than 0.1 were excluded to reduce the number of false polymorphic loci due to sequencing error. The resulting SNPs were exported in GenePop format for further analysis. Since missing data can influence genetic clustering and diversity estimates, we applied three different thresholds for missing data: 40%, 20%, and 10%, to compare for congruence across analyses. Filtering was performed using dartR v2.9.2 (Gruber et al. 2018) to exclude loci and individuals with missing data greater than the specified thresholds.

### 2.4 Genotype cluster analysis

To examine patterns of genetic clustering of SNP genotypes, we used two clustering approaches: multivariate analysis and model-based assignment methods. Multivariate analysis was performed using a Principal Component Analysis (PCA) implemented in *dartR::gl.pcoa*. A model-based assignment method implemented in the software ADMIXTURE v1.3.0 (Alexander et al. 2009) was used to infer the number of genetic clusters *K* and estimate individual ancestry coefficients (*q*) using a maximum likelihood approach. The optimal *K* was identified based on the lowest cross-validation error estimated across 10 independent runs (for *K* = 1 to 10). Individual ancestry coefficients (*q*) were then assessed at the optimal *K* with 100 independent runs. Results were examined using ‘pophelper’ v2.3.1 (Francis 2017). Individuals were assigned to a SNP genotype cluster when *q* > 0.9950 corresponding to operationally ‘pure’ individuals and identified as admixed if otherwise (Caniglia et al. 2020).

We also visualized genetic relationships among SNP genotypes by calculating a Euclidean distance matrix among individuals using the *dartR::gl.dist.ind* function and performing hierarchical clustering using the *hclust* function in the R package stats v4.3.1 (R Core Team 2023). A dendrogram was generated using dendextend v1.17.1 (Galili 2015).

To compare genotype cluster assignments between SNP and microsatellite (SSR) markers, published SSR genotype data (https://doi.org/10.5281/zenodo.8273196) was re-analyzed for 55 samples genotyped at 6 SSR loci and with matching SNP genotypes. PCA performed using the R package *adegenet* v2.1.3 (Jombart 2008) revealed three emergent SSR genotype clusters (SSR-1, SSR-2, SSR-3; Figure S1). A Bayesian model-based assignment approach implemented in the software STRUCTURE v.2.3.4 (Pritchard et al. 2000) was then used to estimate individual ancestry coefficients (*q*), at *K = 3* to assign individuals to genotype clusters revealed by PCA. Ten replicate MCMC simulations were performed for each *K* value using an admixture model with correlated allele frequencies. Each run was carried out for 1x10^6^ iterations with an initial burn-in of 100,000 steps. Individuals were assigned to an SSR cluster based on a threshold value of *q* >= 0.9 (Vaha and Primmer 2006), while individuals having *q* values between 0.10 and 0.9 were categorized as admixed (SSR-A; Table S1).

### 2.5 Genetic diversity and differentiation

Genetic differentiation estimators were calculated for SSR and SNP genotype clusters. For SSR genotypes, measures of genetic differentiation between genotype clusters were estimated using *FST* (Weir and Cockerham 1984) and a standardized *GST* (*G’ST*; Hedrick 2005) corrected for highly polymorphic SSR loci, calculated using the package ‘diveRsity’ v.1.9.9 (Keenan et al. 2013). The significance of *FST* and *G’ST* (null hypothesis of genetic homogeneity, Ho: *FST* = 0) was evaluated by estimating the bootstrapped 95% confidence interval (95% CI). For SNP genotypes, we used the dartR package to calculate Weir and Cockerham’s *FST* and associated p-values.

### 2.6 Outlier loci analysis and functional annotation

To identify SNP loci underlying divergence between genotype clusters, outlier loci were identified using three approaches: BayeScan v2.1 (Foll and Gaggiotti 2008), Arlequin v3.5.2.1 (Excoffier and Lischer 2010) and pcadapt v4.3.5 (Luu et al. 2017). BayeScan and Arlequin are Fst-based methods which identify loci that exhibit higher genetic differentiation among defined groups than expected under a neutral model (Ahrens et al. 2018). BayeScan analysis was performed using 20 pilot runs, each consisting of 5,000 iterations, followed by 100,000 iterations with a burn-in of 50,000 iterations. We used a posterior odds (PO) threshold > 10 and q-value < 0.05 after correction for false discovery rate (FDR; Benjamini and Hochberg 1995). Arlequin meanwhile accounts for hierarchical genetic structure in parameter estimations, and outlier analysis was performed using 200,000 simulations and 100 demes with default expected heterozygosity settings. In contrast, pcadapt identifies outliers as loci which contribute significantly to population structure following a principal component analysis (Luu et al. 2017). We used an alpha value of 0.05 and used the package qvalue v2.1.12 (Storey et al. 2024) to calculate FDR corrected q-values, setting a threshold of q < 0.05 for outlier loci identification. All loci identified by any of the three methods were considered as candidate outliers.

To characterize the outlier loci, contigs were queried against NCBI non-redundant database using BLASTx v 2.15.0 (Altschul et al., 1990). BLAST search was restricted to Echinodermata with an e-value cutoff of 10^−5^. Gene ontology (GO) annotation terms of the outlier loci were retrieved from Gene Ontology database (Gene Ontology Consortium 2023) through UniProt (UniProt Consortium 2023). GO terms were summarized using REVIGO (Supek et al. 2011), a clustering algorithm that relies on semantic similarity measures, and were clustered using the simRel score for functional similarity, allowing for redundancy in similar terms up to a value of 0.7 before removal.

## 3 Results

### 3.1 ddRADseq data filtering and SNP genotyping

A total of 644,788,232 raw reads consisting of forward reads (146 bp) and reverse reads (151 bp) were obtained from 96 individuals collected from three sites (STA, MAB, PUJ). Eleven of the 96 samples were excluded due to low read recovery accounting for only 0.13% (892,096) of the total reads. Quality control and contaminant filtering yielded 459,329,688 reads consisting of sequences from 85 samples: 40 from STA, 21 from MAB and 24 from PUJ, with a total of 1,941,591 variable sites and 4,296,347 heterozygous SNPs identified among individuals. The depth of coverage per individual ranged from 9x – 23x with an average read depth of 14x across all samples. A total of 9,788 SNPs were identified from the remaining 85 individuals. Three individuals with high percentage of missing data (> 60% missing data) were excluded before filtering was conducted at varying thresholds of missing data. Applying three filtering thresholds for maximum percentage of missing data across loci and individual genotypes yielded the following datasets (Table S2): (1) 40% (9788 loci, 82 individuals), (2) 30% (5011 loci, 80 individuals) and (3) 10% (2410 loci, 75 individuals). Results from the 40% missing dataset are presented, unless indicated otherwise, based on concordance of results across the three SNP filtering thresholds.

### 3.2 Genotype cluster analysis

Cluster analysis of SNP genotypes using various approaches consistently recovered three groups that broadly corresponded with mitochondrial lineage and microsatellite genotype clusters. ADMIXTURE analysis identified the optimal number of clusters at *K* = 3 across all datasets with varying missing data thresholds. The majority of the samples were assigned to one of three clusters, hereafter designated as clusters SA-1, SA-2 and SA-3, based on a threshold of *q* > 0.9950. A small number of samples exhibited mixed ancestry (0.0050 < *q* < 0.9950) and were identified as admixed (SA-A; Table 2, Figure 2a), with the number of admixed individuals varying across missing data thresholds (n = 4 to 7, Table S3). For admixed individuals, the highest *q*-values ranged from *q* = 0.800 to *q* = 0.974. Assignment to clusters was broadly concordant across all missing data thresholds, except for three samples identified as parental or pure categories at 40% missing data but identified as admixed at 20% and 10% missing data (Figure S2; Table S3). All admixed individuals occurred in one population (STA) and have ancestry from clusters SA-1 or SA-3 (Figure 2a).

**Figure 2.**
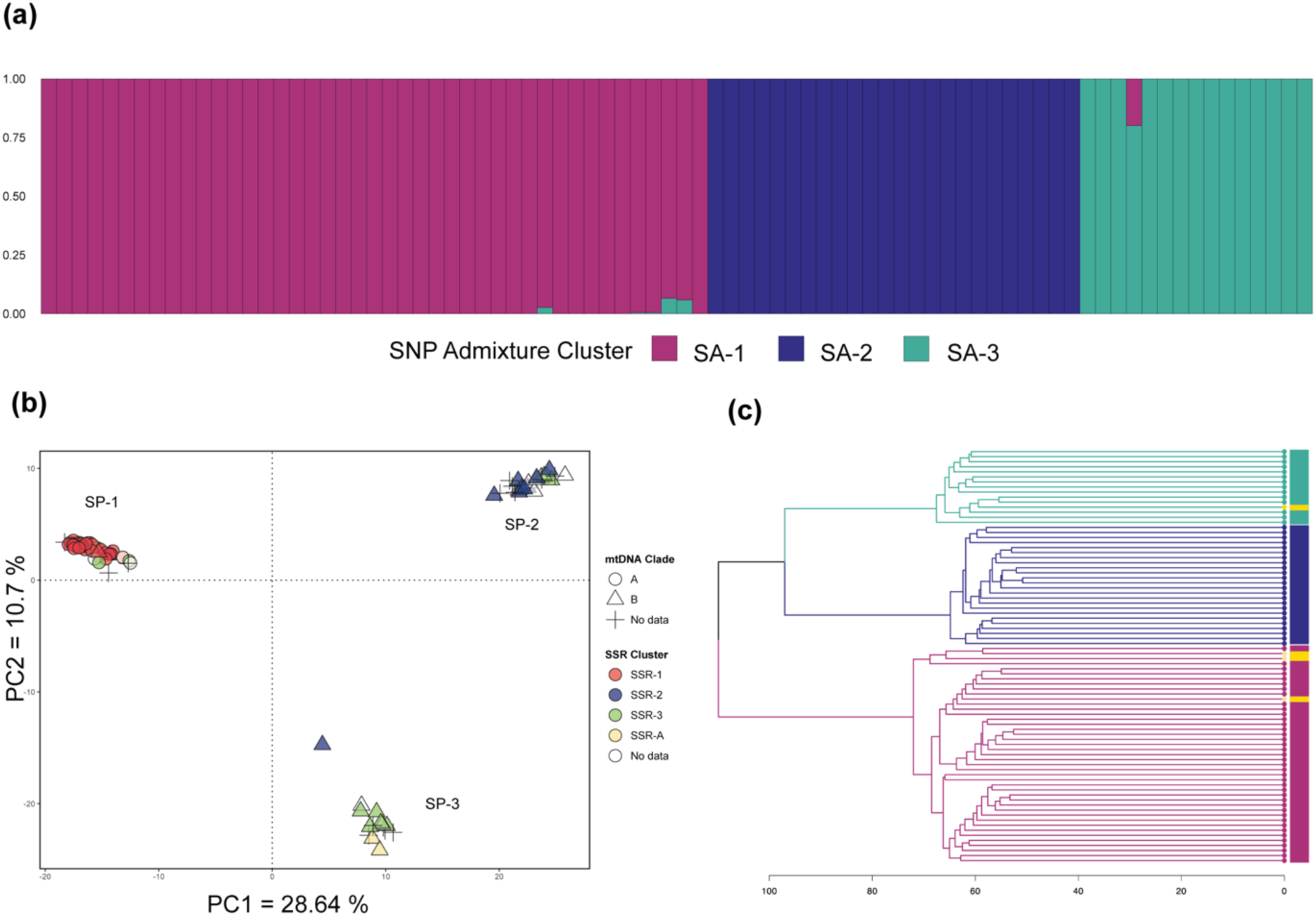
Cluster analysis of *Stichopus* cf. *horrens* SNP genotypes. (a) Barplot of individual ancestry coefficients (*q*) from ADMIXTURE analysis of 82 *Stichopus* cf. *horrens* samples genotyped at 9,788 SNP loci. Each bar on the x-axis represents an individual, the y-axis is the proportion of ancestry in each of three identified clusters, *K*: SA-1, SA-2, SA-3; (b) Principal component analysis of the same individuals showing three genetic lineages (SP-1, SP-2, and SP-3). Each point represents an individual, with lineages identified by shape (mtDNA Clade) and color (SSR genotype clusters). Individuals that were not typed at mtDNA and SSRs (ND = no data) are indicated as crosses (+); (c) Dendrogram of the same individuals based on genetic distance. Lineages are colored according to ADMIXTURE assignments (colored bars beside the dendrogram), with admixed individuals colored yellow.

**Table 1.** Sample information for *Stichopus* cf. *horrens* analyzed in this study. Sample location includes collection site (Location, Site Code), georeference (Latitude, Lat; Longitude, Long). The number of individuals genotyped for mitochondrial COI lineage (Clade A, Clade B), microsatellites (SSR), SNP loci (SNP), and assigned to three SNP lineages (SA-1, SA-2, SA-3) and admixed individuals (SA-A) are shown.

| Location | Site Code | Lat (°N) | Long (°E) | mtDNA |  | SSR | SNP | SNP Clusters |  |  |  |
| --- | --- | --- | --- | --- | --- | --- | --- | --- | --- | --- | --- |
|  |  |  |  | Clade A | Clade B |  |  | SA-1 | SA-2 | SA-3 | SA-A |
| Sta. Ana, Cagayan | STA | 18.511 | 122.152 | 4 | 22 | 17 | 37 | 2 | 17 | 14 | 4 |
| Mabini, Davao de Oro | MAB | 7.297 | 125.836 | 16 | 3 | 18 | 21 | 20 | 1 |  |  |
| Pujada Bay, Davao Oriental | PUJ | 6.844 | 126.228 | 17 | 7 | 20 | 24 | 18 | 6 |  |  |
| **TOTAL** |  |  |  | 37 | 32 | 55 | 82 | 40 | 24 | 14 | 4 |

**Table 2.**
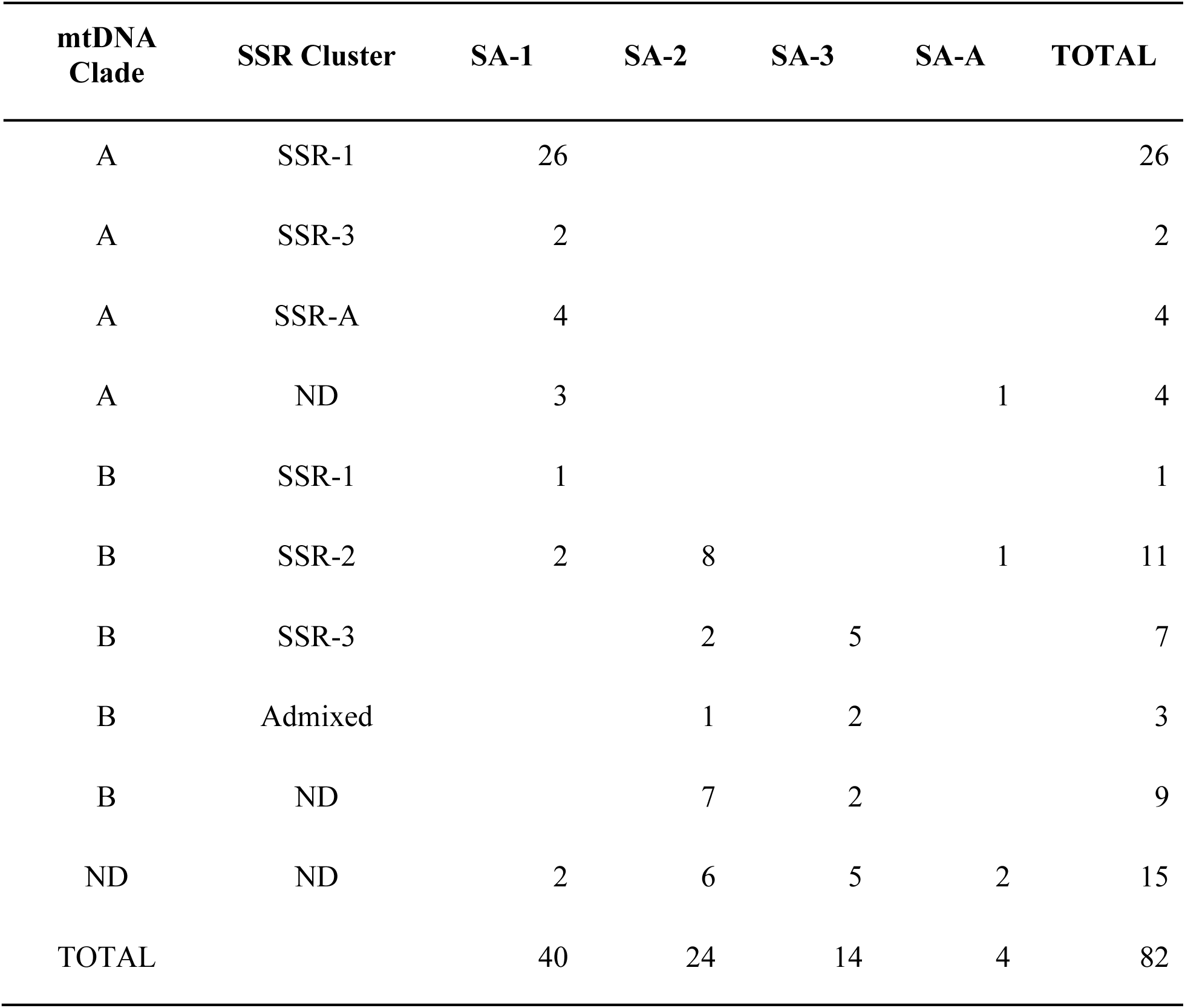
Count of *Stichopus* cf. *horrens* individual assignments to two 1 mtCOI lineages (mtDNA Clade), three microsatellite clusters (SSR Cluster), and three SNP genotype clusters (SA-1, 3 SA-2, SA-3) and individuals with mixed nuclear marker ancestry for microsatellite and SNP 4 loci (SSR-A, SA-A). ND = No data available.

| mtDNA<br>Clade | SSR Cluster | SA-1 | SA-2 | SA-3 | SA-A | TOTAL |
| --- | --- | --- | --- | --- | --- | --- |
| A | SSR-1 | 26 |  |  |  | 26 |
| A | SSR-3 | 2 |  |  |  | 2 |
| A | SSR-A | 4 |  |  |  | 4 |
| A | ND | 3 |  |  | 1 | 4 |
| B | SSR-1 | 1 |  |  |  | 1 |
| B | SSR-2 | 2 | 8 |  | 1 | 11 |
| B | SSR-3 |  | 2 | 5 |  | 7 |
| B | Admixed |  | 1 | 2 |  | 3 |
| B | ND |  | 7 | 2 |  | 9 |
| ND | ND | 2 | 6 | 5 | 2 | 15 |
| TOTAL |  | 40 | 24 | 14 | 4 | 82 |

The PCA plot segregates samples into three distinct, non-overlapping groups, designated as SP-1, SP-2, and SP-3 (Figure 2b). Excluding admixed individuals (SA-A), cluster assignments were congruent between PCA and ADMIXTURE, i.e. SP-1 = SA-1, SP-2 = SA-2, and SP-3 = SA-3 (Table S4). The primary axis (PC1 = 28.64% of the total variance), separated the 82 samples into two groups broadly concordant with mitochondrial lineages: one group consists predominantly of Clade A individuals (SP-1; n = 34 of 38 samples), while the second group consists exclusively of Clade B individuals (SP-2, SP-3; n = 27). The second PCA axis (PC2 = 10.7% of the total variance) further segregates Clade B individuals into two groups broadly corresponding to microsatellite genotype clusters (SSR Cluster), with SP-2 consisting mostly of SSR Cluster 2 individuals (n = 11 of 18 individuals), and SP-3 consisting mostly of SSR Cluster 3 individuals (n = 5 of 9 individuals).

Hierarchical clustering of pairwise individual genetic distances likewise reveals three clusters broadly concordant with PCA and ADMIXTURE assignments (Figure 2c). Clusters SA-2 and SA-3 are genetically more similar, with a smaller mean pairwise Euclidean distance (mean = 88.7, range = 78.9 – 97), than either SA-2 or SA-3 is to SA-3 (SA-1 vs SA-2, mean = 98.0, range = 82.6 – 110.0; SA-1 vs SA-3, mean = 92.3, range 78.2 – 101). Genotype clusters SA-1 and SA-2 occurred in all three sites, with SA-1 being more abundant than SA-2 (48.7% and 29.2% of samples, respectively). Cluster SA-3, accounting for 17% of the samples, was restricted to STA.

### 3.3 Genetic differentiation

The three SNP clusters exhibit significant genetic differentiation. Overall *FST* (global *FST* = 0.458, p < 0.001, n = 78 excluding admixed individuals) and pairwise comparisons between SNP clusters indicate significant differentiation (*FST* > 0): SA-1 and SA-2 (*FST* = 0.510), SA-1 and SA-3 (*FST* = 0.434), SA-2 and SA-3 (*FST* = 0.423). Differentiation between clusters is maintained even in sympatry, i.e. in PUJ (SA1 and SA-2, *FST* = 0.5057, p < 0.001) and STA (SA-2 and SA-3, *FST* = 0.422, p < 0.001), and exhibits much higher *FST* values compared to within-cluster comparisons for different sites. Cluster SA-1 PUJ and MAB populations and Cluster SA-2 PUJ and STA populations exhibit much lower *FST* values than sympatric clusters (*FST* = 0.015 and *FST* = 0.004; p < 0.001; Figure S3a), despite their geographic separation (200 km and 1,400 km, respectively).

The three microsatellite clusters likewise exhibit genetic differentiation (overall *FST* = 0.177, *G’ST* = 0.591). All pairwise *FST* comparisons reject the null hypothesis of homogeneity (*FST* > 0): SSR Cluster 1 and SSR Cluster 2 (*FST* = 0.226), SSR Cluster 2 and SSR Cluster 3 (*FST* = 0.035), SSR Cluster 1 and SSR Cluster 3 (*FST* = 0.156). Differentiation between clusters is significant even in sympatry in MAB (SSR Cluster 1 and Cluster 2, *FST* = 0.231, p < 0.001) and STA (Cluster 2 and Cluster 3 STA, *FST* = 0.176, p < 0.001; Figure S3b). SSR Cluster 2 STA and MAB populations exhibit lower *FST* values (*FST* = 0.156) compared to sympatric lineages at these sites. SSR Cluster 1 populations in MAB and PUJ are not significantly differentiated (*FST* = 0), in contrast with differentiation detected by SNP markers.

### 3.4 Comparison of lineage assignments across mitochondrial, microsatellite, and SNP markers

Comparing lineage assignments across multiple marker types reveals broad concordance of SNP clusters with mtDNA and SSR lineage assignments, for 54 individuals with data across all three marker types (Table 2, Figure 3). While mtDNA lineages recover only two lineages, these correspond with SNP clusters SA-1 (Clade A; n = 32 of 35 SA-1 individuals), SA-2-and SA-3 (Clade B; all 18 SA-2 and SA-3 individuals). Meanwhile, individual assignments based on PCA and STRUCTURE analysis of SSR genotypes at *K* = 3 groups generally correspond with SNP clusters (Table 2): SA-1 consists mostly of SSR-1 (n = 26 of 35 individuals), SA-2 is mostly SSR- 2 (n = 8 of 11 individuals), and SA-3 is predominantly SSR-3 (n = 5 of 8 individuals).

**Figure 3.**
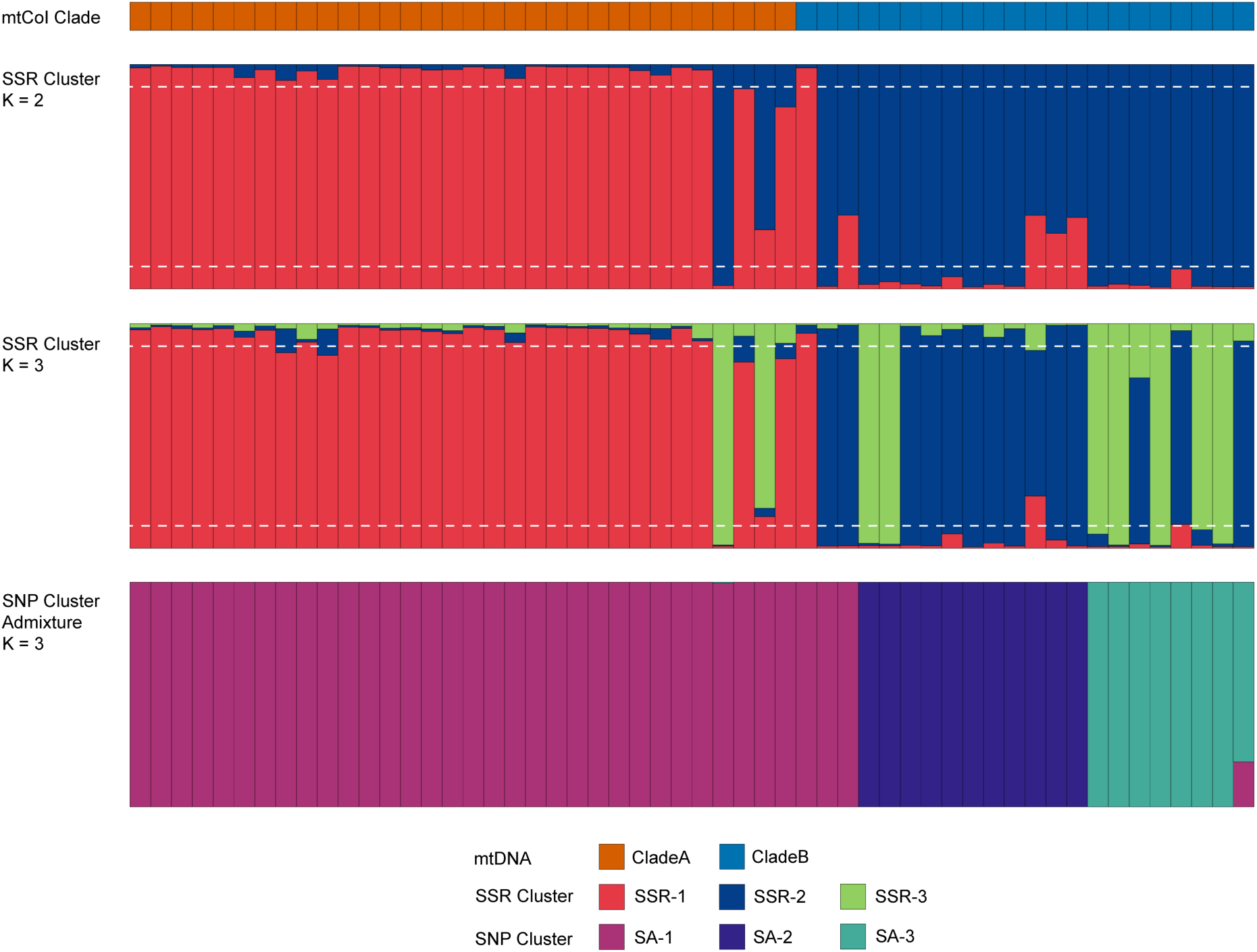
Comparison of lineage assignment barplots for 54 *Stichopus* cf. *horrens* individuals for mtDNA, SSR (at K =2 and K = 3), and SNP markers. Each individual is represented by a vertical bar where the proportion of ancestry (*q*) in a cluster is indicated by colors corresponding to mitochondrial lineages (Clade A, Clade B), microsatellite genotype clusters inferred from STRUCTURE analysis at K = 2 and K = 3, and SNP genotype clusters inferred from ADMIXTURE analysis at K = 3. The horizontal dashed line represents *q-*value thresholds for identifying admixed individuals for SSR data (0.1 < *q* < 0.9).

Excluding one SNP genotype with mixed ancestry (SA-A), discordant assignments were observed between mtDNA and SNP lineages in three of 53 samples (SA-1 genotypes with Clade B lineage), and between SSR and SNP lineages in 13 of 53 samples. Majority of the mismatched SSR cluster assignments were identified as admixed by SSRs, but not by SNPs (8 of 13 assignments), while the remaining mismatches were accounted for by SA-1 identified as SSR Cluster 2 or SSR Cluster 3 (n = 4), and SA-3 identified as SSR Cluster 2 (n = 2) (Table 2, Figure 4).

**Figure 4.**
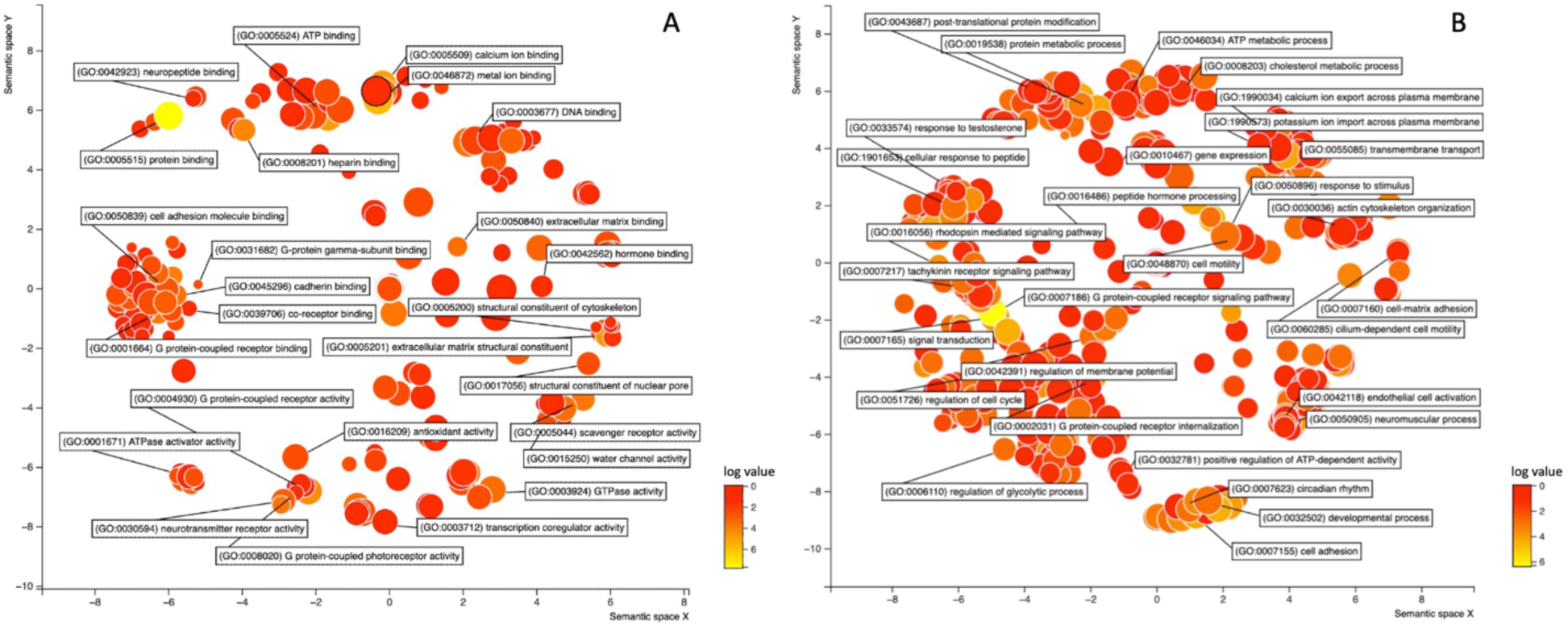
The scatterplot shows the cluster representatives of GO terms (A. molecular function, B. biological process) remaining after the redundancy reduction in a two-dimensional space derived by applying multidimensional scaling to a matrix of the GO terms’ semantic similarities. Bubble color indicates the frequency of the GO term provided as input and bubble size indicates the frequency of the GO term in the underlying GO database (bubbles of more general terms are larger).

### 3.5 SNP outlier analysis and annotation

A total of 108 outlier SNP loci were identified by BayeScan with statistically significant patterns of genetic differentiation. In addition, a total of 384 outlier loci were detected by Arlequin and 1162 outlier loci were detected by pcadapt (Figure S4). All 1507 loci detected using the three methods were considered candidate loci potentially under selection and were subjected to annotation. A total of 157 outlier loci were successfully annotated from the pool of outliers detected with an e-value cutoff of 10^-5^ and were mostly mapped against *Apostichopus japonicus and Holothuria leucospilota*. Functional annotation revealed GO terms related to cell-matrix adhesion, response to testosterone, neuromuscular process, transmembrane transport, peptide hormone binding and receptor signaling (Figure 4).

## 4 Discussion

The present study provides genomic evidence of further cryptic species within *Stichopus* cf. *horrens* and confirms the existence of an additional cryptic species within the Clade B lineage of *Stichopus* cf. *horrens,* first reported by Lizano et al. (2024). Using high-resolution SNP markers, we confirm the presence of three divergent lineages within *Stichopus* cf. *horrens* that maintain reproductive isolation despite occurring in sympatry. These SNP genotype lineages are broadly concordant with mitochondrial lineage and microsatellite genotype assignments. We also identify several outlier loci underlying genomic divergence which provide insight into putative genes contributing to the development and maintenance of reproductive barriers among sympatrically-occuring cryptic lineages of *Stichopus* cf*. horrens*.

### 4.1 SNP loci reveal reproductive isolation among *Stichopus* cf. *horrens* lineages

Using double-digest RAD sequencing to interrogate genetic divergence among previously described *Stichopus* cf. *horrens* lineages provided unparalleled resolution for identifying genetic clusters, with 9,788 SNP loci recovering three distinct groups (SA-1, SA-2, and SA-3). Individual assignments were consistent across various clustering approaches and were also relatively robust to the proportion of missing data. While a small proportion of individuals (3 of 82 genotypes) exhibited discordant assignments across three missing data thresholds, this involved identification of pure individuals as admixed or vice-versa, and there were no mis- assignments of individuals between pure clusters.

Genetic differentiation of sympatric SNP clusters, coupled with broad concordance with mitochondrial and SSR lineages, provides further evidence for reproductive isolation. Genetic differentiation among *S.* cf. *horrens* SNP lineages (*FST* = 0.458) is similar to values for other cryptic echinoderm species also delineated using SNP markers, such as *Stronglyocentrotus* (*FST* range = 0.467-0.497, Addison and Kim 2018) and *Ophioderma* (*FST* range = 0.191 - 0.472; Weber et al. 2019). SNP-based *FST* estimates are also higher than microsatellite-based *FST* values, indicating greater resolution of SNP loci at detecting genetic divergence (Morin et al. 2004). Previous studies suggest that SNP allele lineages are less prone to homoplasy than microsatellites (Coates et al. 2009), and in the case of this study, the small number of microsatellite markers likely accounts for weaker resolving power (Osborne et al., 2022). The low proportion of admixed SNP genotypes (4 of 82 samples) point to limited contemporary gene flow. Moreover, ancestry coefficients of admixed individuals (maximum *q* value range = 0.8 - 0.97) are consistent with older-generation hybrids, and are not typical of recent-generation hybrids, i.e. F1 or F2 where *q* values are expected to be between 0.35 - 0.70 (Caniglia et al. 2020). Thus, these results strongly suggest that the three SNP lineages of *Stichopus* cf. *horrens* represent different species.

### 4.2 Outlier analysis reveal SNP loci related to reproductive processes

Fertilization in broadcast-spawning invertebrates progresses through a series of interactions between sperm and egg, including sperm chemotaxis, which is the activation and attraction of sperm mediated by chemoattractants released by the egg, acrosome reaction, penetration, and membrane fusion (Vacquier 1998). We detected outlier loci with putative functions related to the fertilization process in broadcast spawning invertebrates. Outlier loci detected had relevant BLAST hits with proteins encompassing different functions such as rhodopsin and tachykinin receptor signaling, neurotransmitter receptor activity, response to testosterone, peptide hormone processing, cell-matrix adhesion, sperm head and sperm motility. Most of the outlier loci detected have functions involved in regulating the reproductive process by acting as receptors or by regulating receptor activity. High differentiation at these regions suggests the potential role of these outlier loci in the formation of pre-zygotic barriers among divergent lineages of *Stichopus* cf. *horrens*.

Several outliers identified in this study are involved in cell signaling, a key process in cellular development and its basic machinery involves a receptor that perceives signals such as light, hormones, or neurotransmitters (Trewavas & Malho 1997). Many of these outliers mapped to G protein-coupled receptors (GPCRs), known for their roles in chemosensory reception. Although GPCRs are functionally pleiotropic, previous studies have shown how they can negatively or positively regulate reproduction, i.e. cannabinoid receptors (Rhodopsin type GPCR) found on sea urchin sperm are receptors of endocannabinoids which prevent polyspermy by blocking the acrosome reaction (Chang et al., 1993; Schuel et al., 1994) or binding of gonadotropic neuropeptides to GPCRs to induce spawning (Iwakoshi et al., 1995; Kato et al., 2009; Ohtani et al., 1999, Fujiwara et al., 2010; Yamano et al., 2013). In addition, putative tachykinin-like receptors are GPCRs found to stimulate sperm motility (Satake et al. 2013) and are also identified to be involved with hormone reception in zebrafish (Biran et al. 2012). The outlier loci detected in in this study, suggests the potential role of chemosensory signaling in spawning and previous studies have already shown how hormonal reception influences spawning behaviour in sea cucumbers (Hamel & Mercier, 1999; Kato et al., 2009; Marquet et al. 2018).

Aggregation and synchrony in spawning plays a huge part in the reproductive success of broadcast spawners by offsetting sperm dilution due to water currents (Pennington 1985; Levitan 2000) and is the mechanism by which sympatric Hawaiian limpet species have diverged (Bird et al., 2011). *Stichopus* cf. *horrens* are known to display aggregative behavior during spawning and *in situ* spawning observations have shown that Clade A and Clade B individuals spawn at different times. Clade A individuals spawned 3-4 days before the new moon, between 22:00 and 02:00h of the next day while Clade B individuals spawned 1-4 days after the new moon, at an earlier time, between 19:00 and 23:00h (Juinio-Meñez et al., 2024). How chemosensory reception influences the synchrony of spawning in *Stichopus* cf. *horrens* is unknown. However, earlier studies may provide valuable evidence on how chemical communication affects aggregation and synchrony in spawning of sea cucumbers. While photoperiod is a major trigger of gametogenesis in marine invertebrates (Pearse & Cameron, 1991; McClintock & Watts, 1990; Nelson et al. 2010), chemical communication plays a key role in fine tuning these processes in broadcast spawning invertebrates (Hamel & Mercier, 1996; Soong et al. 2005; Mercier & Hamel, 2010; Marquet et al. 2018). Mercier and Hamel (1996) have shown that gametogenesis was highly asynchronous in *Cucumaria frondosa* individuals kept separately even under natural conditions (Hamel & Mercier, 1996) and that the mucus secreted by *C. frondosa* was found to induce gametogenic synchrony (Hamel & Mercier, 1999). *Holothuria arguinensis* male sea cucumbers were also found to release chemicals that induces aggregation and spawning (Marquet et al. 2018). Male-first spawning behaviour is common among echinoderms, and it is generally observed that male sea cucumbers are more likely to respond to environmental cues than females (Mercier & Hamel, 2010). Therefore, one possible explanation would be that males are triggered to spawn in response to certain environmental variables such as lunar phase which will subsequently synchronize spawning using biological cues such as pheromones. *Stichopus* cf. *horrens* males spawn first during a specific lunar phase followed afterwards by the spawning of females (Juinio-Meñez et al., 2024). This is similar with the previous observation in the crown-of-thorns starfish where males are found to be more responsive to environmental cues and subsequently synchronizes spawning by using pheromones to induce spawning in females and other males (Caballes and Pratchett 2017). Divergence in pheromone receptors may thus exhibit variation in response to cues and ultimately in synchronicity and spawning success. Currently, there is no direct evidence on how divergence in hormone receptors can form a reproductive barrier in sea cucumbers but a previous study on brittle stars found compelling evidence of positive selection on sperm-expressed genes that encode the sodium–proton exchanger (NHE) and the tetrameric potassium-selective cyclic nucleotide-gated channel (TetraKCNG). These two genes form part of the signal transduction cascade within the sperm in response to pheromones (Weber et al., 2017). Moreover, numerous other studies in chemosensory systems have likewise shown how chemosensory receptors can be key players in speciation (Smadja and Butlin 2009; van Schooten et al., 2020).

In echinoderms, where visual capabilities are largely confined to photoreceptors (Millot 1976, Yamamoto et al., 1978, Blevins and Johnsen 2004), and predator avoidance, localization of food sources and species recognition is mostly mediated through chemical signaling (McClintock and Vernon, 1990; Campbell et al. 2001; Morishita and Baretto 2011, Lawrence 2013), the role of chemical receptors in survival is undeniably important. Variation in receptors vital to reproduction may thus affect synchrony of broadcast spawning species and temporal differences in gamete release is one of the primary mechanisms contributing to prezygotic reproductive isolation between closely related marine species in sympatry (Palumbi 1994). The widespread occurrence of synchronous spawning among marine organisms suggests that this trait is strongly favored by natural selection. However, among potentially hybridizing species in sympatry and where hybrids have lower fitness than parental lineages, selection may thus favor asynchrony in gamete release (Geyer and Palumbi 2003, Wolstenholme, J. K 2004, Fogarty et al 2012, Monteiro et al, 2016). The evolution of pre-zygotic isolating mechanisms to limit hybridization is crucial in maintaining species cohesion and may be key factors driving speciation in sympatry (Palumbi 1994; Gardner 1997; Fukami et al. 2003; Levitan et al. 2004; Coyne and Orr, 2004). How divergence in putative hormone receptors can facilitate aggregation and spawning synchrony in sympatric species of *Stichopus* cf. *horrens* remains to be uncovered.

### 4.3 Implications for species identification

Mitochondrial and microsatellite markers can diagnose two lineages, Clade A-Cluster1 and Clade B-Cluster 2, which matches gross morphological differences in papillae distribution and density, but not spicule morphology (Lizano et al. 2024) and is further characterized by asynchronous spawning (Juinio-Meñez et al 2024). Genome-wide SNP markers provide improved resolution over mitochondrial and microsatellite markers, clearly recovering a third lineage, previously detected as a sub-lineage within *Stichopus* cf. *horrens* Clade B-Cluster 2.

Apart from the genetic data reported here, there is limited information on the morphology, ecology, and reproductive biology of the third cryptic species. These results call for a more comprehensive reassessment of the morphology, genetic variation, and ecology of *Stichopus* cf. *horrens* across its distributional range. The SNP markers reported in this study represent novel genomic resources that can be leveraged to develop sequence-based methods for species identification which can accelerate species assessments and monitoring efforts to support broader-scale studies on the biology, ecology and taxonomy of this species complex.

## 5 Conclusions

Findings from this study provide strong evidence that the three genotype clusters of *Stichopus* cf. *horrens* recovered using genomic SNPs are reproductively isolated and represent cryptic species. The recovery of the third genotype cluster thus confirms the existence of the third species within the Clade B lineage. Further investigation into the morphology, ecology, and reproductive biology of the third cryptic species, as well as broader reassessments of *Stichopus* cf. *horrens* across its distribution range are still needed. *FST* outlier analysis from this study revealed a set of highly divergent SNP loci that are mapped to putative genes involved in reproductive processes, such as G-protein coupled receptors (GPCRs). The detection of outlier

SNPs in gene regions related to receptor signaling and hormone response and where a growing body of evidence points to hormone receptors as key players to spawning synchrony (Hamel & Mercier, 1999; Mercier & Hamel, 2010; Marquet *et al*. 2018) suggests that these pathways may play a key role in the reproductive isolation in sympatric populations of *Stichopus* cf. *horrens.* Findings from the present work provide a basis for further examination of the role of divergence in hormone receptors in the formation of pre-zygotic isolating mechanism among sympatric individuals with external fertilization.

## Data Accessibility Statement

All raw sequence files are deposited in NCBI with the BioProject accession number: PRJNA1168564. SNP data and associated metadata are deposited in Zenodo 10.5281/zenodo.13887755.

## Author Contributions

KMKim: Conceptualization (equal), Data curation, Formal analysis (lead), Investigation, Writing – original draft, Writing – review & editing; ADLizano: methodology- sample collection, DNA extraction, Writing – review & editing; RJToonen: resources, Writing – review & editing; RRGotanco: Conceptualization (equal), Formal analysis, Funding acquisition, Project administration, Supervision, Writing – original draft, Writing – review & editing.

### Acknowledgements

This work was supported by the Department of Science and Technology – Philippine Council for Agriculture, Aquatic and Natural Resources Research and Development (DOST- PCAARRD. The authors are grateful to the following individuals who facilitated sample collection and provided information and observations on Stichopus ecology and distribution: Marie Antonette Juinio-Meñez, Rose Angeli Rioja (UP MSI), Ruberto Alforque (BFAR- Romblon), Corazon Batoy (Holy Name University), Helen Bangi (Cagayan State University), Nonita Cabacaba (BFAR Guiuan), Christine Edullantes, Girley Gumanao (Davao del Norte State College), Nadia Palomar-Abesamis (Silliman University), Abduraji S. Tahil (MSU Tawi- Tawi). The authors also thank Faith Paran and Deo Macahig for their assistance with sample collection.

## Ethical Approval

No ethical considerations to declare

## Notes

### Competing Interest Statement

The authors have declared no competing interest.

