## Supplementary Figures for "Genomic divergence of sympatric lineages within *Stichopus* cf. *horrens* (Echinodermata: Stichopodidae): Insights on reproductive isolation inferred from SNP markers"

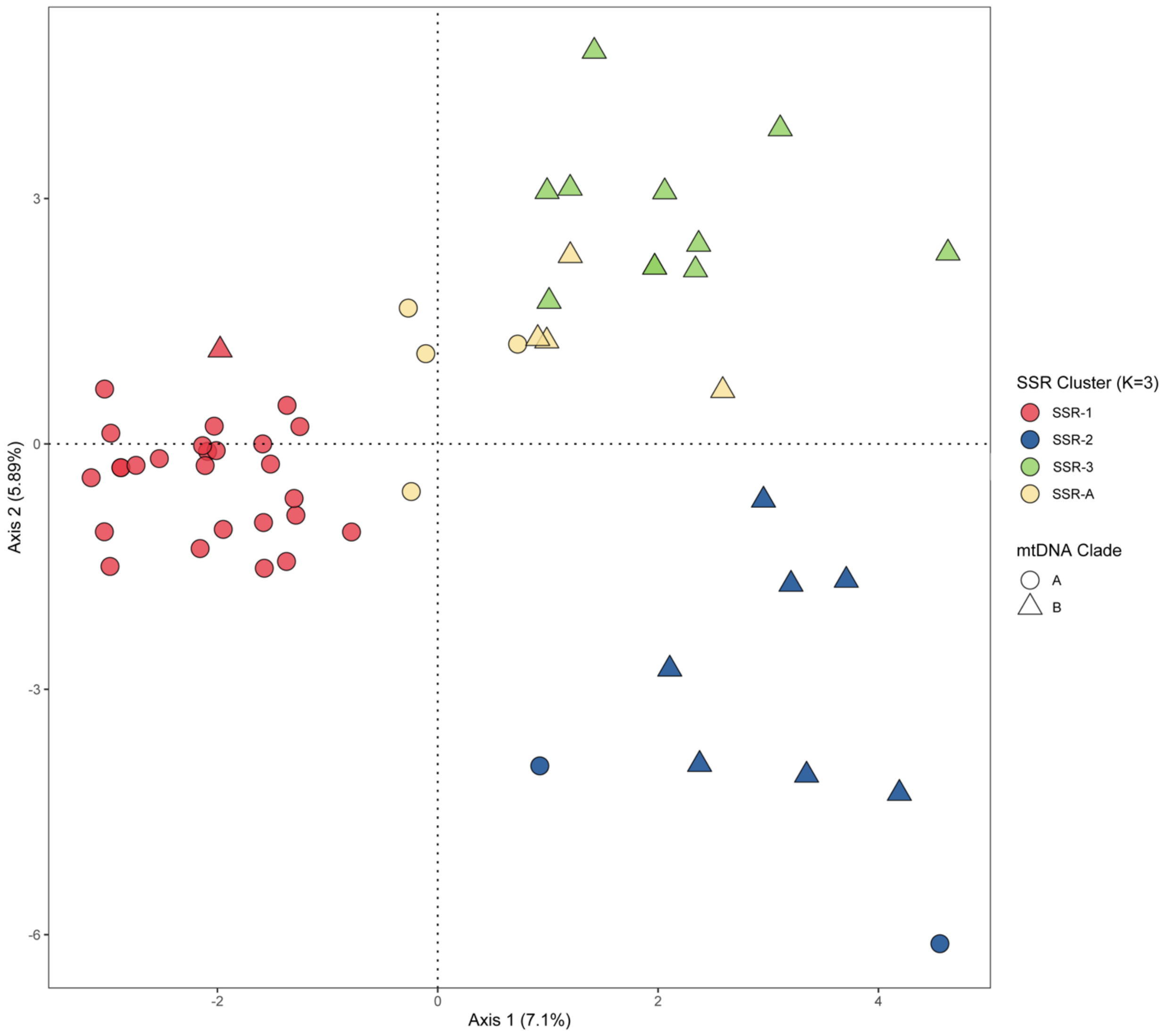


Figure S1. Principal component analysis plot of microsatellite genotypes from 55 *Stichopus* cf. *horrens*. Each point represents an individual, with lineages identified by shape (mtDNA Clade) and color (microsatellite genotype clusters or SSRs). SSR clusters were inferred from STRUCTURE analysis at $K = 3$ groups, with individuals assigned to a cluster when the ancestry coefficient ($q$) is > 0.9, and admixed if $0.1 < q < 0.9$.

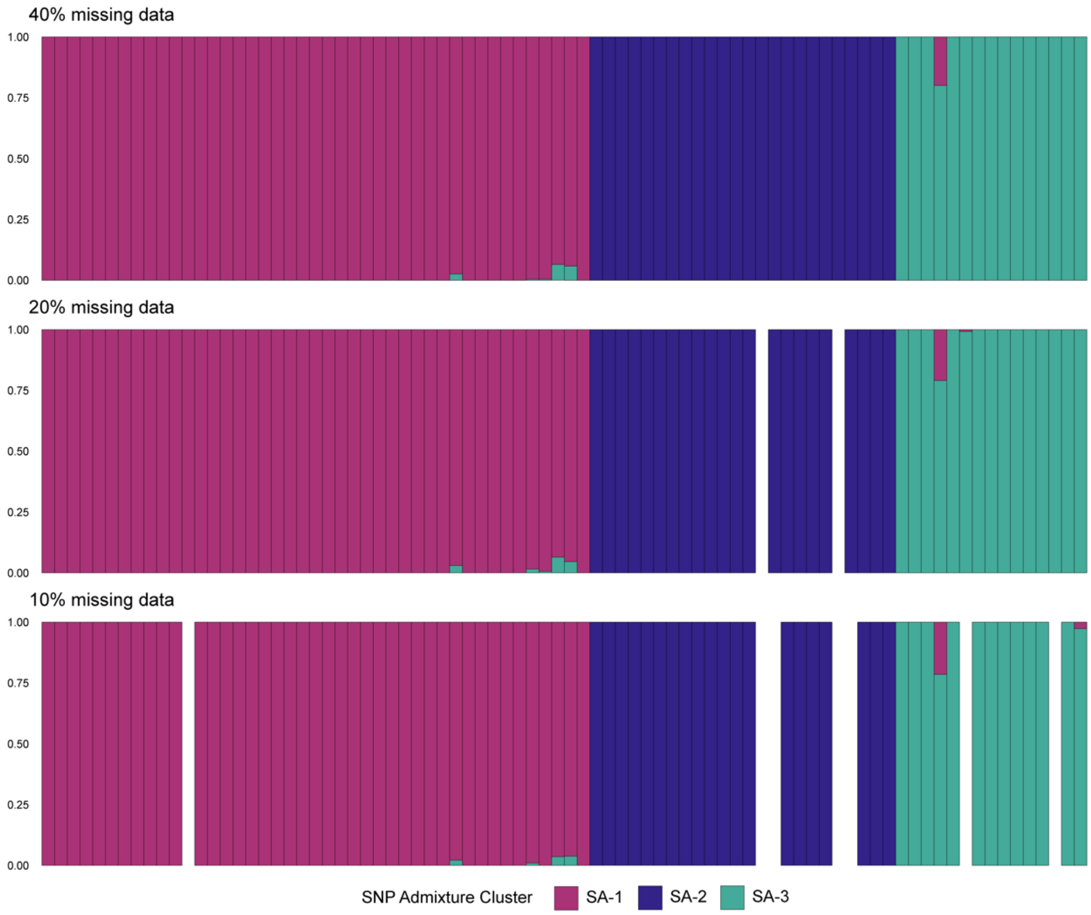


Figure S2. Bar plots of individual ancestry based on Admixture analysis at $K = 3$ genetic clusters across three SNP (Single Nucleotide Polymorphism) datasets with varying levels of missing data (40%, 20%, 10% missing data). Each bar represents one individual, with bar color representing the proportion of ancestry ($q$) in each genetic cluster. White bars represent individuals which were excluded after filtering for missing data.

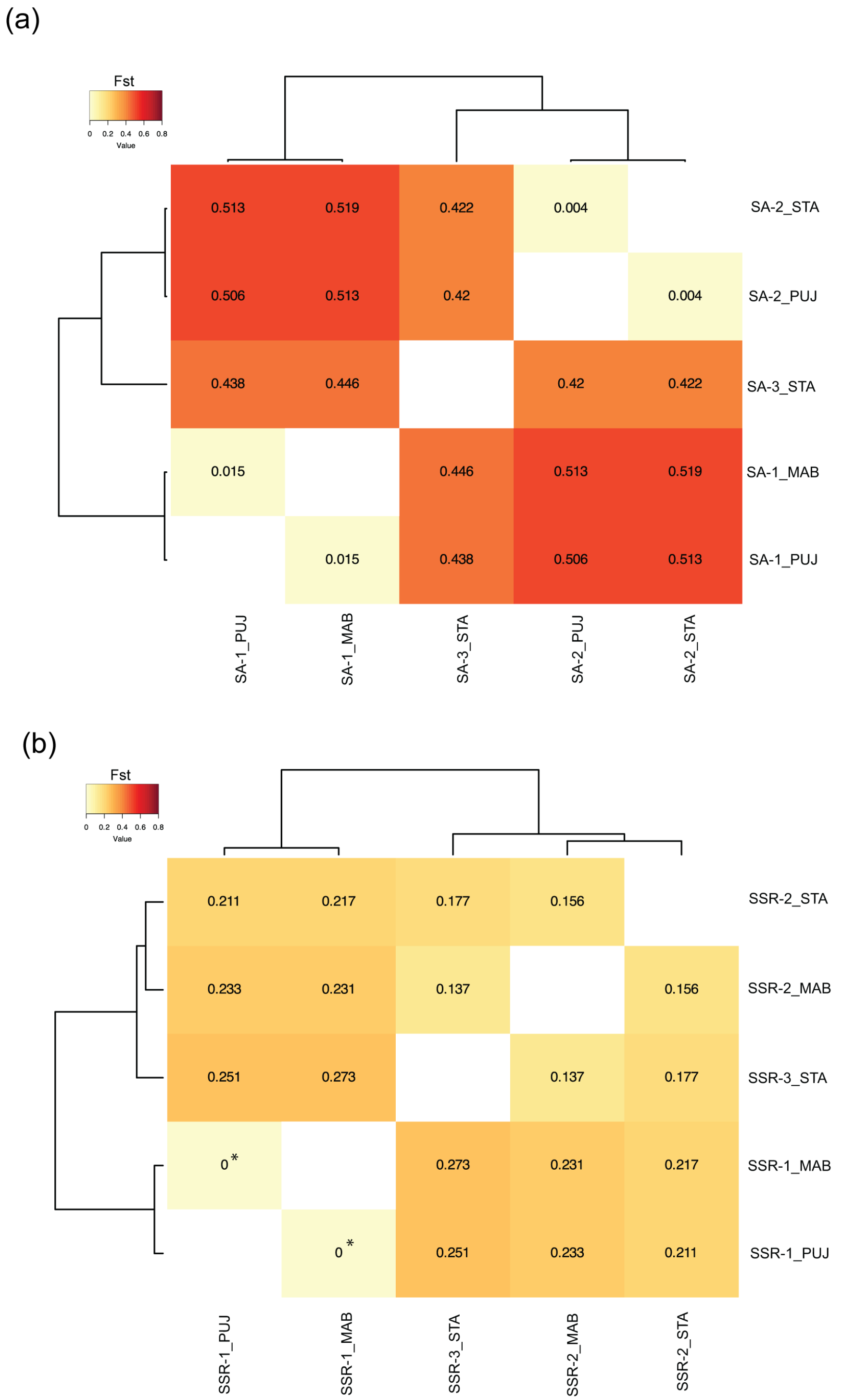


Figure S3. Heatmap of pairwise $F_{ST}$ values for *Stichopus* cf. *horrens* lineages at three collection sites. (a) SNP lineages for 78 individuals based on 9,788 SNP loci. (b) SSR lineages for 44 individuals based on 6 microsatellite loci. Groups with less than five colonies were excluded from the analysis. Hierarchical clustering is represented by the dendrogram, showing three genetic lineages. Non-significant $F_{ST}$ values ($F_{ST} = 0$) are marked with an asterisk.

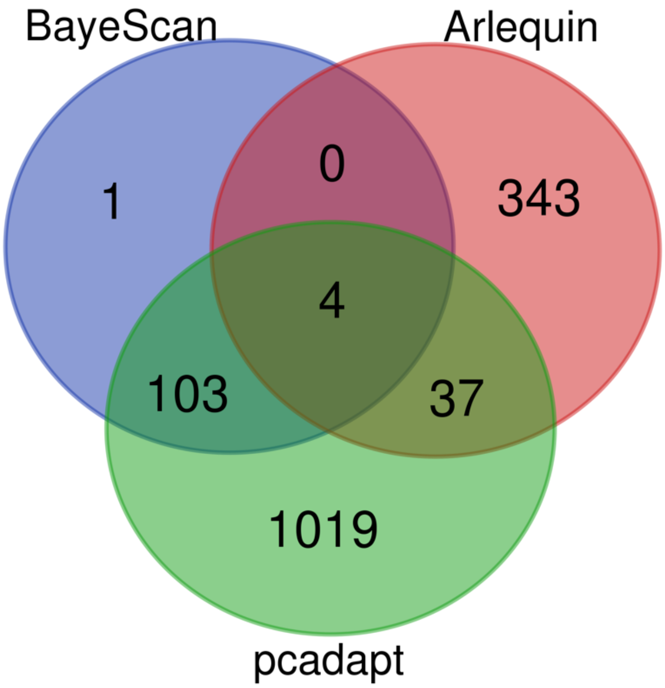


Figure S4. Venn diagram showing the number of outlier SNPs identified across different outlier detection methods and the overlap between these methods.
