## Supplementary Tables for "Genomic divergence of sympatric lineages within *Stichopus* cf. *horrens* (Echinodermata: Stichopodidae): Insights on reproductive isolation inferred from SNP markers"

Table S1. MtDNA clade, SSR and SNP genotype cluster assignments for *Stichopus* cf. *horrens* specimens used in the study. Lir
mtDNA and SSRs (SSR K = 2) are based on the dataset and analysis from Lizano et al. (2024). Assignments for SSR (K = 3) are ba
from Lizano et al. (2024). Assignments for SNP lineages are from data generated in this study, analyzed using PCA and Admixtur

| | | | | SSR | | SN |
|---|---|---|---|---|---|---|
| **Number** | **Sample** | **Site Code** | **mtDNA Clade** | **SSR K = 2** | **SSR K = 3** | **SNP Cluster (PCA)** |
| 1 | STA_15_04 | STA | A | - | - | SP-1 |
| 2 | STA_15_11 | STA | A | - | - | SP-1 |
| 3 | STA_15_48 | STA | A | P2 | SSR-3 | SP-1 |
| 4 | STA_17_20 | STA | - | - | - | SP-1 |
| 5 | STA_17_21 | STA | - | - | - | SP-1 |
| 6 | STA_15_38 | STA | B | - | - | SP-2 |
| 7 | STA_15_06 | STA | B | - | - | SP-2 |
| 8 | STA_15_07 | STA | B | - | - | SP-2 |
| 9 | STA_15_18 | STA | B | P2 | SSR-3 | SP-2 |
| 10 | STA_15_19 | STA | B | P2 | SSR-3 | SP-2 |
| 11 | STA_15_32 | STA | B | P2 | SSR-2 | SP-2 |
| 12 | STA_15_34 | STA | B | P2 | SSR-2 | SP-2 |
| 13 | STA_15_39 | STA | B | P2 | SSR-2 | SP-2 |
| 14 | STA_15_43 | STA | B | P2 | SSR-2 | SP-2 |
| 15 | STA_15_47 | STA | B | P2 | SSR-2 | SP-2 |
| 16 | STA_15_54 | STA | B | - | - | SP-2 |
| 17 | STA_15_58 | STA | - | - | - | SP-2 |
| 18 | STA_17_14 | STA | - | - | - | SP-2 |
| 19 | STA_17_03 | STA | - | - | - | SP-2 |
| 20 | STA_17_08 | STA | - | - | - | SP-2 |
| 21 | STA_17_01 | STA | - | - | - | SP-2 |
| 22 | STA_17_11 | STA | - | - | - | SP-2 |

| | | | | | | |
|---|---|---|---|---|---|---|
| 23 | STA_17_10 | STA | - | - | - | SP-3 |
| 24 | STA_17_15 | STA | - | - | - | SP-3 |
| 25 | STA_15_36 | STA | B | P2 | SSR-3 | SP-3 |
| 26 | STA_15_50 | STA | B | P2 | SSR-2 | SP-3 |
| 27 | STA_17_07 | STA | - | - | - | SP-3 |
| 28 | STA_15_23 | STA | B | - | - | SP-3 |
| 29 | STA_15_24 | STA | B | - | - | SP-3 |
| 30 | STA_15_26 | STA | B | A | SSR-2 | SP-3 |
| 31 | STA_15_35 | STA | B | P2 | SSR-A | SP-3 |
| 32 | STA_15_37 | STA | B | P2 | SSR-3 | SP-3 |
| 33 | STA_15_40 | STA | B | P2 | SSR-A | SP-3 |
| 34 | STA_15_41 | STA | - | - | - | SP-3 |
| 35 | STA_15_44 | STA | B | P2 | SSR-3 | SP-3 |
| 36 | STA_15_45 | STA | B | P2 | SSR-3 | SP-3 |
| 37 | STA_17_05 | STA | - | - | - | SP-3 |
| 38 | MAB_16_11 | MAB | A | P1 | SSR-1 | SP-1 |
| 39 | MAB_16_12 | MAB | A | P1 | SSR-1 | SP-1 |
| 40 | MAB_16_13 | MAB | A | P1 | SSR-1 | SP-1 |
| 41 | MAB_16_14 | MAB | A | P1 | SSR-1 | SP-1 |
| 42 | MAB_16_17 | MAB | B | P2 | SSR-2 | SP-1 |
| 43 | MAB_16_18 | MAB | A | A | SSR-A | SP-1 |
| 44 | MAB_16_02 | MAB | A | P1 | SSR-1 | SP-1 |
| 45 | MAB_16_20 | MAB | A | - | - | SP-1 |
| 46 | MAB_16_21 | MAB | A | P1 | SSR-1 | SP-1 |
| 47 | MAB_16_22 | MAB | A | P1 | SSR-1 | SP-1 |
| 48 | MAB_16_25 | MAB | A | P1 | SSR-A | SP-1 |
| 49 | MAB_16_26 | MAB | A | P1 | SSR-1 | SP-1 |
| 50 | MAB_16_28 | MAB | A | P1 | SSR-A | SP-1 |
| 51 | MAB_16_03 | MAB | A | P1 | SSR-1 | SP-1 |
| 52 | MAB_16_30 | MAB | B | A | SSR-2 | SP-1 |
| 53 | MAB_16_31 | MAB | - | - | - | SP-1 |

| | | | | | | |
|---|---|---|---|---|---|---|
| 54 | MAB_16_32 | MAB | - | - | - | SP-1 |
| 55 | MAB_16_04 | MAB | A | P1 | SSR-1 | SP-1 |
| 56 | MAB_16_06 | MAB | A | P1 | SSR-1 | SP-1 |
| 57 | MAB_16_08 | MAB | A | P1 | SSR-1 | SP-1 |
| 58 | MAB_16_07 | MAB | B | P2 | SSR-2 | SP-2 |
| 59 | PUJ_16_13 | PUJ | A | P1 | SSR-1 | SP-1 |
| 60 | PUJ_16_14 | PUJ | A | - | - | SP-1 |
| 61 | PUJ_16_16 | PUJ | A | P1 | SSR-1 | SP-1 |
| 62 | PUJ_16_17 | PUJ | A | P1 | SSR-1 | SP-1 |
| 63 | PUJ_16_18 | PUJ | A | A | SSR-3 | SP-1 |
| 64 | PUJ_16_19 | PUJ | A | A | SSR-A | SP-1 |
| 65 | PUJ_16_23 | PUJ | A | P1 | SSR-1 | SP-1 |
| 66 | PUJ_16_05 | PUJ | A | P1 | SSR-1 | SP-1 |
| 67 | PUJ_16_06 | PUJ | A | P1 | SSR-1 | SP-1 |
| 68 | PUJ_16_07 | PUJ | A | P1 | SSR-1 | SP-1 |
| 69 | PUJ_16_08 | PUJ | A | P1 | SSR-1 | SP-1 |
| 70 | PUJ_16_09 | PUJ | A | P1 | SSR-1 | SP-1 |
| 71 | PUJ_16_21 | PUJ | A | P1 | SSR-1 | SP-1 |
| 72 | PUJ_16_24 | PUJ | A | P1 | SSR-1 | SP-1 |
| 73 | PUJ_16_26 | PUJ | A | P1 | SSR-1 | SP-1 |
| 74 | PUJ_16_28 | PUJ | A | P1 | SSR-1 | SP-1 |
| 75 | PUJ_16_29 | PUJ | A | P1 | SSR-1 | SP-1 |
| 76 | PUJ_16_22 | PUJ | B | P1 | SSR-1 | SP-1 |
| 77 | PUJ_16_02 | PUJ | B | A | SSR-A | SP-2 |
| 78 | PUJ_16_10 | PUJ | B | - | - | SP-2 |
| 79 | PUJ_16_11 | PUJ | B | A | SSR-2 | SP-2 |
| 80 | PUJ_16_12 | PUJ | B | A | SSR-2 | SP-2 |
| 81 | PUJ_16_15 | PUJ | B | - | - | SP-2 |
| 82 | PUJ_16_20 | PUJ | B | - | - | SP-2 |

ıeage assignments for
sed on reanalysis of data
re approaches.

| IP |
| --- |
| **SNP Cluster (Admixture)** |
| SA-A |
| SA-1 |
| SA-1 |
| SA-A |
| SA-A |
| SA-2 |
| SA-2 |
| SA-2 |
| SA-2 |
| SA-2 |
| SA-2 |
| SA-2 |
| SA-2 |
| SA-2 |
| SA-2 |
| SA-2 |
| SA-2 |
| SA-2 |
| SA-2 |
| SA-2 |
| SA-2 |
| SA-2 |

| |
|---|
| SA-3 |
| SA-3 |
| SA-3 |
| SA-A |
| SA-3 |
| SA-3 |
| SA-3 |
| SA-3 |
| SA-3 |
| SA-3 |
| SA-3 |
| SA-3 |
| SA-3 |
| SA-3 |
| SA-3 |
| SA-1 |
| SA-1 |
| SA-1 |
| SA-1 |
| SA-1 |
| SA-1 |
| SA-1 |
| SA-1 |
| SA-1 |
| SA-1 |
| SA-1 |
| SA-1 |
| SA-1 |
| SA-1 |
| SA-1 |
| SA-1 |

| |
|---|
| SA-1 |
| SA-1 |
| SA-1 |
| SA-1 |
| SA-2 |
| SA-1 |
| SA-1 |
| SA-1 |
| SA-1 |
| SA-1 |
| SA-1 |
| SA-1 |
| SA-1 |
| SA-1 |
| SA-1 |
| SA-1 |
| SA-1 |
| SA-1 |
| SA-1 |
| SA-1 |
| SA-1 |
| SA-1 |
| SA-1 |
| SA-2 |
| SA-2 |
| SA-2 |
| SA-2 |
| SA-2 |
| SA-2 |

Table S2. Information on three SNP datasets following SNP filtering at three levels of missing dat
missing) across loci and genotypes.

| Filtering Strategy | % missing loci | % missing genotypes | N loci | |
|---|---|---|---|---|
| | | | | STA |
| Unfiltered | 40 | | 9788 | 40 |
| Relaxed | 40 | 40 | 9788 | 37 |
| Moderate | 20 | 20 | 5011 | 37 |
| Strict | 10 | 10 | 2410 | 35 |

:a thresholds (40%, 3020% and 10%

| N individuals | | |
| --- | --- | --- |
| MAB | PUJ | Total |
| 21 | 24 | 85 |
| 21 | 24 | 82 |
| 21 | 24 | 80 |
| 20 | 24 | 75 |

Table S3. Individual ancestry coefficients (q) and SNP lineages (SNP Cluster) based on ADMIXTU
were assigned to three lineages (SA-1, SA-2, SA-3) when q>0.995, and identified as admixed (SA-

| Number | Sample | Site Code | 40% missing data | | | 20 |
|---|---|---|---|---|---|---|
| | | | q1 | q2 | q3 | q1 |
| 1 | STA_15_04 | STA | 0.97413 | 0.00001 | 0.02586 | 0.97003 |
| 2 | STA_15_11 | STA | 0.99560 | 0.00001 | 0.00439 | 0.98386 |
| 3 | STA_15_48 | STA | 0.99565 | 0.00001 | 0.00434 | 0.99351 |
| 4 | STA_17_20 | STA | 0.93494 | 0.00001 | 0.06505 | 0.93536 |
| 5 | STA_17_21 | STA | 0.94203 | 0.00001 | 0.05796 | 0.95380 |
| 6 | STA_15_38 | STA | 0.00001 | 0.99998 | 0.00001 | 0.00001 |
| 7 | STA_15_06 | STA | 0.00001 | 0.99998 | 0.00001 | 0.00001 |
| 8 | STA_15_07 | STA | 0.00001 | 0.99998 | 0.00001 | 0.00001 |
| 9 | STA_15_18 | STA | 0.00001 | 0.99998 | 0.00001 | 0.00001 |
| 10 | STA_15_19 | STA | 0.00001 | 0.99998 | 0.00001 | 0.00001 |
| 11 | STA_15_32 | STA | 0.00001 | 0.99998 | 0.00001 | 0.00001 |
| 12 | STA_15_34 | STA | 0.00001 | 0.99998 | 0.00001 | - |
| 13 | STA_15_39 | STA | 0.00001 | 0.99998 | 0.00001 | 0.00001 |
| 14 | STA_15_43 | STA | 0.00001 | 0.99998 | 0.00001 | 0.00001 |
| 15 | STA_15_47 | STA | 0.00001 | 0.99998 | 0.00001 | 0.00001 |
| 16 | STA_15_54 | STA | 0.00001 | 0.99998 | 0.00001 | 0.00001 |
| 17 | STA_15_58 | STA | 0.00001 | 0.99998 | 0.00001 | 0.00001 |
| 18 | STA_17_14 | STA | 0.00001 | 0.99998 | 0.00001 | - |
| 19 | STA_17_03 | STA | 0.00001 | 0.99998 | 0.00001 | 0.00001 |
| 20 | STA_17_08 | STA | 0.00001 | 0.99998 | 0.00001 | 0.00001 |
| 21 | STA_17_01 | STA | 0.00001 | 0.99998 | 0.00001 | 0.00001 |
| 22 | STA_17_11 | STA | 0.00001 | 0.99998 | 0.00001 | 0.00001 |
| 23 | STA_17_10 | STA | 0.00001 | 0.00001 | 0.99998 | 0.00001 |
| 24 | STA_17_15 | STA | 0.00001 | 0.00001 | 0.99998 | 0.00001 |
| 25 | STA_15_36 | STA | 0.00001 | 0.00001 | 0.99998 | 0.00001 |
| 26 | STA_15_50 | STA | 0.19955 | 0.00001 | 0.80044 | 0.20948 |
| 27 | STA_17_07 | STA | 0.00001 | 0.00001 | 0.99998 | 0.00001 |
| 28 | STA_15_23 | STA | 0.00001 | 0.00001 | 0.99998 | 0.00762 |
| 29 | STA_15_24 | STA | 0.00001 | 0.00001 | 0.99998 | 0.00001 |
| 30 | STA_15_26 | STA | 0.00001 | 0.00001 | 0.99998 | 0.00001 |
| 31 | STA_15_35 | STA | 0.00001 | 0.00001 | 0.99998 | 0.00001 |
| 32 | STA_15_37 | STA | 0.00001 | 0.00001 | 0.99998 | 0.00001 |
| 33 | STA_15_40 | STA | 0.00001 | 0.00001 | 0.99998 | 0.00001 |
| 34 | STA_15_41 | STA | 0.00001 | 0.00001 | 0.99998 | 0.00001 |
| 35 | STA_15_44 | STA | 0.00001 | 0.00001 | 0.99998 | 0.00001 |
| 36 | STA_15_45 | STA | 0.00001 | 0.00001 | 0.99998 | 0.00001 |
| 37 | STA_17_05 | STA | 0.00001 | 0.00001 | 0.99998 | 0.00001 |
| 38 | MAB_16_11 | MAB | 0.99998 | 0.00001 | 0.00001 | 0.99998 |
| 39 | MAB_16_12 | MAB | 0.99998 | 0.00001 | 0.00001 | 0.99998 |
| 40 | MAB_16_13 | MAB | 0.99998 | 0.00001 | 0.00001 | 0.99998 |
| 41 | MAB_16_14 | MAB | 0.99998 | 0.00001 | 0.00001 | 0.99998 |

| | | | | | | |
|---|---|---|---|---|---|---|
| 42 | MAB_16_17 | MAB | 0.99998 | 0.00001 | 0.00001 | 0.99998 |
| 43 | MAB_16_18 | MAB | 0.99998 | 0.00001 | 0.00001 | 0.99998 |
| 44 | MAB_16_02 | MAB | 0.99998 | 0.00001 | 0.00001 | 0.99998 |
| 45 | MAB_16_20 | MAB | 0.99998 | 0.00001 | 0.00001 | 0.99998 |
| 46 | MAB_16_21 | MAB | 0.99998 | 0.00001 | 0.00001 | 0.99998 |
| 47 | MAB_16_22 | MAB | 0.99998 | 0.00001 | 0.00001 | 0.99998 |
| 48 | MAB_16_25 | MAB | 0.99998 | 0.00001 | 0.00001 | 0.99998 |
| 49 | MAB_16_26 | MAB | 0.99998 | 0.00001 | 0.00001 | 0.99998 |
| 50 | MAB_16_28 | MAB | 0.99998 | 0.00001 | 0.00001 | 0.99998 |
| 51 | MAB_16_03 | MAB | 0.99998 | 0.00001 | 0.00001 | 0.99998 |
| 52 | MAB_16_30 | MAB | 0.99998 | 0.00001 | 0.00001 | 0.99998 |
| 53 | MAB_16_31 | MAB | 0.99998 | 0.00001 | 0.00001 | 0.99998 |
| 54 | MAB_16_32 | MAB | 0.99998 | 0.00001 | 0.00001 | 0.99998 |
| 55 | MAB_16_04 | MAB | 0.99998 | 0.00001 | 0.00001 | 0.99998 |
| 56 | MAB_16_06 | MAB | 0.99998 | 0.00001 | 0.00001 | 0.99998 |
| 57 | MAB_16_08 | MAB | 0.99998 | 0.00001 | 0.00001 | 0.99998 |
| 58 | MAB_16_07 | MAB | 0.00001 | 0.99998 | 0.00001 | 0.00001 |
| 59 | PUJ_16_13 | PUJ | 0.99998 | 0.00001 | 0.00001 | 0.99998 |
| 60 | PUJ_16_14 | PUJ | 0.99998 | 0.00001 | 0.00001 | 0.99998 |
| 61 | PUJ_16_16 | PUJ | 0.99998 | 0.00001 | 0.00001 | 0.99998 |
| 62 | PUJ_16_17 | PUJ | 0.99998 | 0.00001 | 0.00001 | 0.99998 |
| 63 | PUJ_16_18 | PUJ | 0.99998 | 0.00001 | 0.00001 | 0.99998 |
| 64 | PUJ_16_19 | PUJ | 0.99998 | 0.00001 | 0.00001 | 0.99998 |
| 65 | PUJ_16_23 | PUJ | 0.99998 | 0.00001 | 0.00001 | 0.99998 |
| 66 | PUJ_16_05 | PUJ | 0.99998 | 0.00001 | 0.00001 | 0.99998 |
| 67 | PUJ_16_06 | PUJ | 0.99998 | 0.00001 | 0.00001 | 0.99998 |
| 68 | PUJ_16_07 | PUJ | 0.99998 | 0.00001 | 0.00001 | 0.99998 |
| 69 | PUJ_16_08 | PUJ | 0.99998 | 0.00001 | 0.00001 | 0.99998 |
| 70 | PUJ_16_09 | PUJ | 0.99998 | 0.00001 | 0.00001 | 0.99998 |
| 71 | PUJ_16_21 | PUJ | 0.99998 | 0.00001 | 0.00001 | 0.99998 |
| 72 | PUJ_16_24 | PUJ | 0.99998 | 0.00001 | 0.00001 | 0.99998 |
| 73 | PUJ_16_26 | PUJ | 0.99998 | 0.00001 | 0.00001 | 0.99998 |
| 74 | PUJ_16_28 | PUJ | 0.99998 | 0.00001 | 0.00001 | 0.99998 |
| 75 | PUJ_16_29 | PUJ | 0.99998 | 0.00001 | 0.00001 | 0.99998 |
| 76 | PUJ_16_22 | PUJ | 0.99998 | 0.00001 | 0.00001 | 0.99998 |
| 77 | PUJ_16_02 | PUJ | 0.00001 | 0.99998 | 0.00001 | 0.00001 |
| 78 | PUJ_16_10 | PUJ | 0.00001 | 0.99998 | 0.00001 | 0.00001 |
| 79 | PUJ_16_11 | PUJ | 0.00001 | 0.99997 | 0.00002 | 0.00001 |
| 80 | PUJ_16_12 | PUJ | 0.00001 | 0.99998 | 0.00001 | 0.00001 |
| 81 | PUJ_16_15 | PUJ | 0.00001 | 0.99998 | 0.00001 | 0.00001 |
| 82 | PUJ_16_20 | PUJ | 0.00001 | 0.99998 | 0.00001 | 0.00001 |

RE analysis of SNP datasets with varying thresholds of missing data (40%, 20%, 10%
-A) otherwise.

| % missing data | | 10% missing data | | | |
|---|---|---|---|---|---|
| q2 | q3 | q1 | q2 | q3 | 40% missing |
| 0.00001 | 0.02997 | 0.97831 | 0.00001 | 0.02169 | SA-A |
| 0.00001 | 0.01613 | 0.98990 | 0.00001 | 0.01010 | SA-1 |
| 0.00001 | 0.00648 | 0.99998 | 0.00001 | 0.00001 | SA-1 |
| 0.00001 | 0.06463 | 0.96371 | 0.00001 | 0.03628 | SA-A |
| 0.00001 | 0.04619 | 0.96204 | 0.00001 | 0.03795 | SA-A |
| 0.99998 | 0.00001 | 0.00001 | 0.99998 | 0.00001 | SA-2 |
| 0.99998 | 0.00001 | 0.00001 | 0.99998 | 0.00001 | SA-2 |
| 0.99998 | 0.00001 | 0.00001 | 0.99998 | 0.00001 | SA-2 |
| 0.99998 | 0.00001 | 0.00001 | 0.99998 | 0.00001 | SA-2 |
| 0.99998 | 0.00001 | 0.00001 | 0.99998 | 0.00001 | SA-2 |
| 0.99998 | 0.00001 | 0.00001 | 0.99998 | 0.00001 | SA-2 |
| - | - | - | - | - | SA-2 |
| 0.99998 | 0.00001 | - | - | - | SA-2 |
| 0.99998 | 0.00001 | 0.00001 | 0.99998 | 0.00001 | SA-2 |
| 0.99998 | 0.00001 | 0.00001 | 0.99998 | 0.00001 | SA-2 |
| 0.99998 | 0.00001 | 0.00001 | 0.99998 | 0.00001 | SA-2 |
| 0.99998 | 0.00001 | 0.00001 | 0.99998 | 0.00001 | SA-2 |
| - | - | - | - | - | SA-2 |
| 0.99998 | 0.00001 | - | - | - | SA-2 |
| 0.99998 | 0.00001 | 0.00001 | 0.99998 | 0.00001 | SA-2 |
| 0.99998 | 0.00001 | 0.00001 | 0.99998 | 0.00001 | SA-2 |
| 0.99998 | 0.00001 | 0.00001 | 0.99998 | 0.00001 | SA-2 |
| 0.00001 | 0.99998 | 0.00001 | 0.00001 | 0.99998 | SA-3 |
| 0.00001 | 0.99998 | 0.00001 | 0.00001 | 0.99998 | SA-3 |
| 0.00001 | 0.99998 | 0.00001 | 0.00001 | 0.99998 | SA-3 |
| 0.00001 | 0.79051 | 0.21487 | 0.00001 | 0.78512 | SA-A |
| 0.00001 | 0.99998 | 0.00001 | 0.00001 | 0.99998 | SA-3 |
| 0.00001 | 0.99237 | - | - | - | SA-3 |
| 0.00001 | 0.99998 | 0.00001 | 0.00001 | 0.99998 | SA-3 |
| 0.00001 | 0.99998 | 0.00001 | 0.00001 | 0.99998 | SA-3 |
| 0.00001 | 0.99998 | 0.00001 | 0.00001 | 0.99998 | SA-3 |
| 0.00001 | 0.99998 | 0.00001 | 0.00001 | 0.99998 | SA-3 |
| 0.00001 | 0.99998 | 0.00001 | 0.00001 | 0.99998 | SA-3 |
| 0.00001 | 0.99998 | 0.00001 | 0.00001 | 0.99998 | SA-3 |
| 0.00001 | 0.99998 | - | - | - | SA-3 |
| 0.00001 | 0.99998 | 0.00001 | 0.00001 | 0.99998 | SA-3 |
| 0.00001 | 0.99998 | 0.02683 | 0.00001 | 0.97316 | SA-3 |
| 0.00001 | 0.00001 | 0.99998 | 0.00001 | 0.00001 | SA-1 |
| 0.00001 | 0.00001 | 0.99998 | 0.00001 | 0.00001 | SA-1 |
| 0.00001 | 0.00001 | 0.99998 | 0.00001 | 0.00001 | SA-1 |
| 0.00001 | 0.00001 | 0.99998 | 0.00001 | 0.00001 | SA-1 |

| | | | | | |
|---|---|---|---|---|---|
| 0.00001 | 0.00001 | 0.99998 | 0.00001 | 0.00001 | SA-1 |
| 0.00001 | 0.00001 | 0.99998 | 0.00001 | 0.00001 | SA-1 |
| 0.00001 | 0.00001 | 0.99998 | 0.00001 | 0.00001 | SA-1 |
| 0.00001 | 0.00001 | 0.99998 | 0.00001 | 0.00001 | SA-1 |
| 0.00001 | 0.00001 | 0.99998 | 0.00001 | 0.00001 | SA-1 |
| 0.00001 | 0.00001 | 0.99998 | 0.00001 | 0.00001 | SA-1 |
| 0.00001 | 0.00001 | 0.99998 | 0.00001 | 0.00001 | SA-1 |
| 0.00001 | 0.00001 | - | - | - | SA-1 |
| 0.00001 | 0.00001 | 0.99998 | 0.00001 | 0.00001 | SA-1 |
| 0.00001 | 0.00001 | 0.99998 | 0.00001 | 0.00001 | SA-1 |
| 0.00001 | 0.00001 | 0.99998 | 0.00001 | 0.00001 | SA-1 |
| 0.00001 | 0.00001 | 0.99998 | 0.00001 | 0.00001 | SA-1 |
| 0.00001 | 0.00001 | 0.99998 | 0.00001 | 0.00001 | SA-1 |
| 0.00001 | 0.00001 | 0.99998 | 0.00001 | 0.00001 | SA-1 |
| 0.00001 | 0.00001 | 0.99998 | 0.00001 | 0.00001 | SA-1 |
| 0.00001 | 0.00001 | 0.99998 | 0.00001 | 0.00001 | SA-1 |
| 0.99998 | 0.00001 | 0.00001 | 0.99998 | 0.00001 | SA-2 |
| 0.00001 | 0.00001 | 0.99998 | 0.00001 | 0.00001 | SA-1 |
| 0.00001 | 0.00001 | 0.99998 | 0.00001 | 0.00001 | SA-1 |
| 0.00001 | 0.00001 | 0.99998 | 0.00001 | 0.00001 | SA-1 |
| 0.00001 | 0.00001 | 0.99998 | 0.00001 | 0.00001 | SA-1 |
| 0.00001 | 0.00001 | 0.99998 | 0.00001 | 0.00001 | SA-1 |
| 0.00001 | 0.00001 | 0.99998 | 0.00001 | 0.00001 | SA-1 |
| 0.00001 | 0.00001 | 0.99998 | 0.00001 | 0.00001 | SA-1 |
| 0.00001 | 0.00001 | 0.99998 | 0.00001 | 0.00001 | SA-1 |
| 0.00001 | 0.00001 | 0.99998 | 0.00001 | 0.00001 | SA-1 |
| 0.00001 | 0.00001 | 0.99998 | 0.00001 | 0.00001 | SA-1 |
| 0.00001 | 0.00001 | 0.99998 | 0.00001 | 0.00001 | SA-1 |
| 0.00001 | 0.00001 | 0.99998 | 0.00001 | 0.00001 | SA-1 |
| 0.00001 | 0.00001 | 0.99998 | 0.00001 | 0.00001 | SA-1 |
| 0.00001 | 0.00001 | 0.99998 | 0.00001 | 0.00001 | SA-1 |
| 0.00001 | 0.00001 | 0.99998 | 0.00001 | 0.00001 | SA-1 |
| 0.00001 | 0.00001 | 0.99998 | 0.00001 | 0.00001 | SA-1 |
| 0.00001 | 0.00001 | 0.99998 | 0.00001 | 0.00001 | SA-1 |
| 0.00001 | 0.00001 | 0.99998 | 0.00001 | 0.00001 | SA-1 |
| 0.99998 | 0.00001 | 0.00001 | 0.99998 | 0.00001 | SA-2 |
| 0.99998 | 0.00001 | 0.00001 | 0.99998 | 0.00001 | SA-2 |
| 0.99998 | 0.00001 | 0.00001 | 0.99998 | 0.00001 | SA-2 |
| 0.99998 | 0.00001 | 0.00001 | 0.99998 | 0.00001 | SA-2 |
| 0.99998 | 0.00001 | 0.00001 | 0.99998 | 0.00001 | SA-2 |
| 0.99998 | 0.00001 | 0.00001 | 0.99998 | 0.00001 | SA-2 |

missing data). Individuals

| SNP Cluster | |
|---|---|
| **20% missing** | **10% missing** |
| SA-A | SA-A |
| SA-A | SA-A |
| SA-A | SA-1 |
| SA-A | SA-A |
| SA-A | SA-A |
| SA-2 | SA-2 |
| SA-2 | SA-2 |
| SA-2 | SA-2 |
| SA-2 | SA-2 |
| SA-2 | SA-2 |
| SA-2 | SA-2 |
| SA-A | SA-A |
| SA-2 | SA-A |
| SA-2 | SA-2 |
| SA-2 | SA-2 |
| SA-2 | SA-2 |
| SA-2 | SA-2 |
| SA-A | SA-A |
| SA-2 | SA-A |
| SA-2 | SA-2 |
| SA-2 | SA-2 |
| SA-2 | SA-2 |
| SA-3 | SA-3 |
| SA-3 | SA-3 |
| SA-3 | SA-3 |
| SA-A | SA-A |
| SA-3 | SA-3 |
| SA-A | SA-A |
| SA-3 | SA-3 |
| SA-3 | SA-3 |
| SA-3 | SA-3 |
| SA-3 | SA-3 |
| SA-3 | SA-3 |
| SA-3 | SA-3 |
| SA-3 | SA-A |
| SA-3 | SA-3 |
| SA-3 | SA-A |
| SA-1 | SA-1 |
| SA-1 | SA-1 |
| SA-1 | SA-1 |
| SA-1 | SA-1 |

| | |
|---|---|
| SA-1 | SA-1 |
| SA-1 | SA-1 |
| SA-1 | SA-1 |
| SA-1 | SA-1 |
| SA-1 | SA-1 |
| SA-1 | SA-1 |
| SA-1 | SA-1 |
| SA-1 | SA-A |
| SA-1 | SA-1 |
| SA-1 | SA-1 |
| SA-1 | SA-1 |
| SA-1 | SA-1 |
| SA-1 | SA-1 |
| SA-1 | SA-1 |
| SA-1 | SA-1 |
| SA-1 | SA-1 |
| SA-2 | SA-2 |
| SA-1 | SA-1 |
| SA-1 | SA-1 |
| SA-1 | SA-1 |
| SA-1 | SA-1 |
| SA-1 | SA-1 |
| SA-1 | SA-1 |
| SA-1 | SA-1 |
| SA-1 | SA-1 |
| SA-1 | SA-1 |
| SA-1 | SA-1 |
| SA-1 | SA-1 |
| SA-1 | SA-1 |
| SA-1 | SA-1 |
| SA-1 | SA-1 |
| SA-1 | SA-1 |
| SA-1 | SA-1 |
| SA-1 | SA-1 |
| SA-1 | SA-1 |
| SA-2 | SA-2 |
| SA-2 | SA-2 |
| SA-2 | SA-2 |
| SA-2 | SA-2 |
| SA-2 | SA-2 |
| SA-2 | SA-2 |

Table S4. Cross-tabulation of SNP cluster assignments from PCA (SP-1, SP-2, SP-3) ar
analyses (SA-1, SA-2, SA-3, SA-A)

| | **SNP Cluster (Admixture)** | | | |
|---|---|---|---|---|
| **SNP Cluster (PCA)** | **SA-1** | **SA-2** | **SA-3** | **SA-A** |
| SP-1 | 40 | 0 | 0 | 3 |
| SP-2 | 0 | 24 | 0 | 0 |
| SP-3 | 0 | 0 | 14 | 1 |
| **TOTAL** | **40** | **24** | **14** | **4** |

ıd ADMIXTURE

| |
|---|
| **TOTAL** |
| 43 |
| 24 |
| 15 |
| **82** |
